# Time-dependent processing of dengue virus polyprotein yields multiple capsid forms that disrupt cellular homeostasis

**DOI:** 10.1101/2025.10.02.679945

**Authors:** Preeti Chauhan, Himanshu Vankhede, Sakshi Rawat, Deepti Maisnam, Lekha Gandhi, Deepika Rathore, Akash Gulyani, Musturi Venkataramana

## Abstract

Dengue virus infections spread to new geographical regions every year, cause significant morbidity and mortality, yet the progress in drug/vaccine development is limited due to an incomplete understanding of its life cycle. While a few of the proteins coded by this virus exist in multiple forms, the sequential events of polyprotein processing to yield them and the consequential effects on host cells are unclear. In this study, the virus infected cell culture data suggest that the polyprotein undergoes a time-dependent processing to yield multiple capsid forms, ie, capsid, capsid-anchor, capsid-anchor-pr, capsid-anchor-prM. Among them, the c-anc and c-anc-pr were found to localize to the mitochondria upon transfection into the HEK cells. The transmission electron microscopy studies suggest that c-anc induces mitochondrial fragmentation. Further studies showed that c-anc affects the mitochondrial fusion-related proteins and the genes involved in their functions. The pulldown experiments indicate that c-anc also interacts with α1-syntrophin (SNTA1), a cytoplasmic protein that plays a role in redox potential, oxidative stress and mitochondrial biogenesis. Importantly, the mitostress analyses suggest that c-anc triggers impaired mitochondrial dysfunctions like potential, respiration and ATP generation. In this study, we also identify ursonic acid (UNA) as a c-anc binding compound that restores mitochondrial functions and suppresses virus multiplication *in vitro*, *ex vivo* and in mice. The revealed disrupted mitochondrial homeostasis appears to be common to DENV, ZIKV, JEV, YFV, HCV, ASFV and SARS-CoV-2; hence, UNA or its related compounds could be considered as inhibitors for the above virus infections.

## Introduction

Dengue virus (DENV) is a mosquito-borne flavivirus endemic in nearly 130 countries. The Center for Disease Control (CDC) estimates that approximately 400 million people suffer from dengue fever each year[1], [2]. Clinically, dengue presents a spectrum of symptoms ranging from asymptomatic or mild febrile illness to life-threatening complications. The majority of DENV infections are either asymptomatic or present as a self-limiting febrile illness with symptoms such as high fever, headache, nausea, rash, and muscle pain[1], [3]. World Health Organization (WHO) classifies dengue symptoms into dengue without warning signs, with warning signs and severe dengue[3]. DENV belongs to the family *Flaviviridae* which also includes West Nile virus (WNV), Yellow fever virus (YFV), Japanese encephalitis virus (JEV) and Tick-borne encephalitis virusL(TBEV)[4]. It is primarily transmitted by *Aedes aegypti* mosquitoes and to a lesser extent by *Aedes albopictus* ^5^. This virus is approximately 50Lnm in diameter with a ∼10.7 kb positive-sense single-stranded RNA genome that encodes a single polyprotein[5], [6]. The polyprotein is co- and post-translationally processed into three structural (capsid, pre-membrane and envelope) and seven non-structural (NS1, NS2A/B, NS3, NS4A/B and NS5) proteins. This processing is mediated by both host and viral proteases[7], [8], [9]. During viral maturation, the polyprotein appears to undergo a series of tightly regulated sequential cleavage events at both the N- and C-termini. Among the structural proteins, capsid (C), membrane (M, derived from prM) and envelope (E) form the main components of the virion. The intermediates, such as the anchor region of the capsid, help to hold the polyprotein to the ER membrane[10], [11]. Of the non-structural proteins, NS3 and NS5 can act independently or in complex with co-factors NS2B and NS4B, respectively[7], [8], [9], [12]. However, the timing, cleavage and regulation of these interactions and their co-expressions remain incompletely understood. Among the virus coded proteins, the capsid protein plays a pivotal role in RNA encapsidation, nucleocapsid formation and virion assembly[13], [14]. The mature DENV capsid is a basic ∼12LkDa homodimer protein with strong affinities for nucleic acids and membranes[13], [15]. Structural studies reveal that it consists of four α-helices per monomer, with a highly charged, unstructured N-terminus and a hydrophobic internal region that facilitates ER membrane binding[13], [16], [17]. Studies have shown that dengue capsid can bind to host core histones and disrupt nucleosomes, potentially interfering with chromatin mediated antiviral responses[18]. It also interacts with host proteins such as nucleolin, influencing both viral replication, host cell homeostasis, and interacts with human Sec3 at the perinuclear and cytoplasmic region[14], [19], [20]. These observations suggest that the capsid’s role extends beyond particle assembly to direct modulation of host cell pathways. Such multifunctionality was also reported in the case of the core protein of hepatitis C virus (HCV) that localizes to mitochondria and modulates both apoptosis and innate immune signalling, while the WNV capsid protein has been shown to induce mitochondrial dysfunction and activate caspases[21], [22], [23], [24]. Although considerable progress has been made in identifying viral targets, there are still no approved antiviral therapies for DENV. Several compounds have shown promise in targeting viral components such as the NS2B/NS3 protease, NS5 polymerase, and even the capsid itself[25], [26], [27], [28]. The recent efforts also highlight host directed approaches, including kinase inhibitors as potential antiviral strategies[29]. However, most of these candidates remain in preclinical stages and progress toward effective dengue therapeutics has been limited.

In this study, the data suggested that the dengue virus polyprotein undergoes time dependent processing to yield multiple forms of capsid protein. Among them, c-anc localizes to the mitochondria, disrupts their homeostasis and function. Ursonic acid was identified as c-anc binding compound and which was found to restore the disrupted mitochondrial homeostasis and functions. It appears that a similar mechanism of capsid mediated disrupted mitochondrial homeostasis is being applied by other viruses like ZIKV, JEV, YFV, ASFV and SARS-CoV2; hence, the ursonic acid or its relevant molecules could be used to treat such infections.

## Materials and Methods

### Cell lines

HEK293 and Vero cells were obtained from the National Centre for Cell Science (NCCS), Pune, India and cultured in Dulbecco’s Modified Eagle Medium (DMEM). K562 cells were maintained in Roswell Park Memorial Institute (RPMI-1640) medium and both were supplemented with 10% FBS. All cells were incubated at 37°C in a humidified atmosphere containing 5% COL. Antibiotic-Antimycotics (Anti-Anti) from Gibco(cat# 15240062) were used at 1% concentration.

### Antibodies

Primary antibodies of MFN1 (Cat#14739), MFN2 (Cat#9482), OPA1 (Cat#80471), DRP1 (Cat#8570), GFP (cat# 2956), SNTA1 (Cat# sc-166634) and Alexa Fluor® 488 (Cat#4412) were from Cell Signalling Technology (CST). All above antibodies were used at 1:2000 dilution. GAPDH (Cat# CAB932Hu01; 1:3000) and anti-rabbit HRP-conjugated secondary antibodies (Cat# sc-2357; 1: 10, 000) were purchased from Santa Cruz Biotechnology, USA. Anti-IFNGR1 (Cat# PA5-96413) and anti-IFNAR1 (Cat# HY-P99137) were purchased from Invitrogen and MedChemExpress, respectively. Anti-NS2BNS3pro and anti-capsid antibodies (1:1000) were in-house raised.

### Plasmids and molecular cloning

All plasmid constructs were generated using a previously characterized DENV-1 strain (KX618706.1) from our laboratory as a template[30]. Viral RNA isolation was performed from 140µl of infected cell culture supernatant using the QiAamp viral RNA isolation kit (Qiagen, Germany). For cDNA synthesis, 1 µg of RNA from each sample was used with 1µl of reverse transcriptase, 5µl of 5x Prime Script buffer (iScript^TM^ cDNA Synthesis Kit) and the final volume was adjusted to 20µl with DEPC-treated water. The reaction mixture was incubated at 25°C for 5 minutes, followed by 46°C for 20 minutes, then at 95°C for 1 minute. Following RT-PCR, the capsid-anchor-pr (c-anc-pr) region was PCR amplified and cloned into pJET 1.2 cloning vector (Thermo Scientific) using T4 DNA ligase and transformed into *E. coli* DH5α cells and confirmed by RT-PCR. For mammalian expression, three regions of capsid gene-capsid (1-100aa), c-anc (1-114aa) and c-anc-pr (1-206aa) were PCR-amplified from the pJET1.2-c-anc-pr using the gene-specific primers, each flanked by *Xho* I (5’) and *Bam* HI (3’) restriction sites. The same forward primer sequence, CGGAAGAAGACGGG was used for all constructs, with restriction enzymes selected based on the respective cloning vectors. The reverse primer sequences were as follows: for capsid, TCTTTTTCTCCTGTTCAT; for capsid anchor, CGCCAGGGCTGTG; and for capsid-anchor-pr, CGCCAGTTTGAGAGC. PCR products and pEGFP-N1 vectors were digested with *Xho* I and *Bam* H I enzymes. The gel eluted fragments were ligated at 37°C for 3 hours to generate in-frame EGFP fusion constructs under the CMV promoter and transformed into *E. coli* DH5α. For bacterial expression, the c-anc region was amplified using primers flanked by *Bam* HI (5’) and *Hind* III (3’) sites. The amplicon and the pET-28a (+) vector were digested with the above enzymes and ligated. The recombinant plasmid was transformed into *E. coli* DH5α. All clones were confirmed by PCR and restriction enzyme digestions.

The nucleotide sequence encoding the SARS-CoV-2 Nucleocapsid (N) protein was PCR-amplified using the N gene specific primer from the pcDNA3.1 SARS-CoV-2 N plasmid (Addgene plasmid #158079, USA) as the template. The amplified gene was digested with 5’*Bam* HI and 3’*Eco* R1 restriction enzymes and ligated to a similarly digested pET-28a(+) vector and transformed into *E. coli* DH5α cells. The colonies were screened using PCR and were confirmed by restriction enzyme digestion.

### Cell transfections and virus infections

HEK293 cells were transfected with pEGFP-N1 vector and plasmids encoding capsid variants; capsid (c), capsid-anchor (c-anc) and capsid-anchor-pr (c-anc-pr) using Lipofectamine 3000 (Invitrogen). Briefly, recombinant plasmid DNA and transfection reagent were mixed in a 1:2 ratio in serum-free DMEM, incubated for 15 minutes at room temperature (RT) and then added dropwise onto the cells. After 5 hours, the transfection medium was replaced with complete DMEM. All downstream analyses were carried out 48 hours post-transfections. For infection studies, cells were seeded one day prior to infection and monitored for confluency. Upon reaching ∼70% confluency, DENV1 infected cell supernatant was added at a multiplicity of infection (MOI) of 0.1. Cells were incubated for 1 hour at 37°C to allow viral adsorption. Then, viral inoculum was removed and media with 2% FBS was added to the cells. All downstream analyses were carried out 72 hours post-infection unless different time points were needed for time dependent experiments.

To examine the conserveness among c-anc proteins of DENV 1-4, the amino acid sequences [DENV1 (ASD49619.1), DENV2 (AYE66919.1), DENV3 (YP_001621843.1), DENV4 (ARB18129.1)] were retrieved from the NCBI protein database. Multiple sequence alignments were performed using Clustal Omega and phylogenetic trees were constructed using the neighbour-joining method and p-distance model in Molecular Evolutionary Genetic Analysis Version 12 (MEGA12)[31].

### Immunofluorescence assay

For transfection-based immunofluorescence, HEK293 cells were seeded on glass coverslips in 12-well plates and transfected with pEGFP-N1 and capsid variants as described above. After 48 hours, cells were washed with 1× phosphate-buffered saline (PBS) and fixed with 4% paraformaldehyde (PFA) for 15 minutes at RT. Nuclei were counterstained with Hoechst 33258 (1:1000 dilution; Thermo Fisher Scientific) and coverslips were mounted using 100% glycerol. Confocal images were captured at 63x magnification.

For infection-based immunofluorescence, the infected Vero cells (as detailed earlier) were washed with PBS and fixed with 4% PFA in PBS for 15 minutes at RT. Cells were then permeabilized with 0.2% Triton X-100 for 5 minutes at RT and blocked with 0.5% (w/v) bovine serum albumin (BSA) in PBS for 1 hour. The cells were incubated with primary antibodies overnight at 4°C, followed by three washes with PBS. Subsequently, cells were incubated with fluorescent conjugated secondary antibodies (Alexa Flour 488) for 2 hours at RT, washed again with PBS and stained with Hoechst 33258. Fluorescence images were captured using a fluorescence microscope at 20x and 60x magnification and the number of positive cells expressing the NS2BNS3 and capsid were recorded at random areas.

### Subcellular fractionation and western blot analysis

The transfected cells were harvested from 100 mm dishes. Total cellular proteins were extracted using RIPA lysis buffer (Sigma-Aldrich, R0278) supplemented with protease inhibitor cocktail (Thermo Fisher Scientific). Incubated on ice for 30Lminutes and centrifuged at 13,000 x g for 20Lminutes at 4°C and the supernatants were collected for immunoblotting. Nuclear fractions were prepared by incubating the PBS washed cells in hypotonic buffer on ice for 10Lmin, followed by centrifugation at 2500 x g for 10Lmin. The nuclear pellet was washed thrice with ice-cold PBS. Mitochondrial fractions were isolated using the mitochondria isolation kit (Abcam; ab110168). Briefly, c-anc transfected cells at a concentration of approximately 5 mg/ml were resuspended in reagent A and incubated on ice for 10 minutes. The cells were then homogenized using a Dounce homogenizer and centrifuged at 1,000 x g for 10 minutes at 4°C. The supernatant was collected and saved. The pellet was resuspended in reagent B, homogenised and spun as above. Both supernatants were mixed and centrifuged again at 10,000 x g for 10 minutes at 4°C. The resulting mitochondrial pellet was resuspended in reagent C and stored in -80°C for further experimental analysis. Protein samples from whole-cell lysates and all subcellular fractions were mixed with 4x SDS PAGE loading buffer and boiled for 5 minutes, resolved on 10% SDS-PAGE and were transferred onto PVDF membranes overnight at 30V at 4°C. After Ponceau S staining, the membranes were blocked in 7% non-fat dry milk dissolved in PBST (PBS + 0.1% Tween-20) for 1hour at RT. The membranes were incubated with respective primary antibodies overnight at 4°C and washed three times (15Lminutes each) in PBST and incubated with HRP-conjugated secondary antibodies for 1hour at RT. Detection was performed using the ChemiDoc MP Imaging System (Bio-Rad) with FemtoLUCENT^TM^ PLUS-HRP (G-Biosciences, Cat# 786-003). For drug treatment analysis, Vero cells were seeded in 6-well plates, pre-treated with UNA (5, 10 and 15LμM) for 6Lhour and infected with dengue virus supernatants. At 72Lh post-infection, cells were lysed in RIPA buffer and the lysates were resolved on 10% SDS-PAGE followed by immunoblotting using anti-NS2BNS3pro and anti-GAPDH antibodies.

### Carbonate extraction and protease protection assays

For the carbonate extraction assay, crude mitochondria were isolated from three 100 mm dishes of pEGFP-c-anc transfected HEK293 cells. Mitochondrial pellet was resuspended in 100 mM NaLCOL (pH ∼11.5), mixed thoroughly and centrifuged at 90,000 x g for 40 minutes at 4°C using a TLA-150 rotor. Supernatant (soluble proteins) and pellet (membrane-associated proteins) fractions were analysed by 10% SDS-PAGE and immunoblot.

For the protease protection assay, the HEK293 cells were transfected with pEGFP-c-anc and after 48 hours the crude mitochondria were isolated. For the proteinase K protection assay, mitochondrial pellets were resuspended in 1× TD buffer (49.99 mM Tris base, 274.13 mM NaCl, 20.12 mM KCl, 14 mM NaLHPOL, pH 7.4) and divided into three equal aliquots. Each aliquot was treated with 100Lμg/mL proteinase K and incubated at RT for 60 minutes[32] The digestion was terminated by the addition of 2LmM PMSF. Samples were analysed by 10% SDS-PAGE and immunoblotted using antibodies against GFP and MFN2 to assess protease accessibility.

### Transmission electron microscopy

HEK293 cells transfected with pEGFP-c-anc and dengue virus infected Vero cells were washed twice with PBS. Then the cells were fixed with 2.5% glutaraldehyde prepared in 0.1M phosphate buffer (pH 7.2) for 24 hours at 4°C, followed by post-fixation treatment for 2 hours in 2% aqueous osmium tetroxide using the same buffer. The cells were undergone a graded ethanol dehydration series, infiltrated and embedded in Araldite 6005 resin. Ultrathin sections (50-70 nm) were made using a glass knife on a Leica Ultracut UCT ultramicrotome. The sections were placed on copper grids, stained with saturated aqueous uranyl acetate, counterstained with Reynolds lead citrate and examined under a JEM-F200 transmission electron microscope (JOEL).

### Transcriptome analysis

Total RNA was extracted from pEGFP-N1 and pEGFP-c-anc transfected HEK293 cells using TRIzol reagent (Thermo Fisher Scientific), following the manufacture’s protocol. RNA sequencing was performed by Eurofins Genomics India Pvt. Ltd. Genes were classified as significantly differentially expressed if they had an adjusted P-value < 0.05, with upregulation indicated by a positive logL fold change and downregulation by a negative logL fold change.

To validate the RNA-seq results, primers were designed for mitochondrial genes ATP6, CO3, CYB, CO2, ND2 and ND6 based on the sequences reported in Wallace 2016[33].

### Expression and purification of recombinant proteins

The constructs containing the c-anc gene was transformed into *E. coli* BL21(DE3) cells, which were cultured in 1 litre of LB medium with 30 µg/mL kanamycin at 37°C with shaking until the OD600 reached 0.6. Protein expression was induced by adding 1mM IPTG and incubating for an additional 4 hours at 37°C. Cells were harvested by centrifugation (8000 rpm, 4°C, 10 min) and the pellet was resuspended in ∼30 mL 1× PBST (pH 7.5) on ice. Cell lysis was performed by sonication, followed by centrifugation (10,000 rpm, 4°C, 50 min) to collect the soluble fraction. The soluble fraction was applied onto the Ni-NTA affinity column (Qiagen, Cat# 30210) pre-equilibrated with lysis buffer (1× PBS, 1% Triton X-100, 2 mM DTT, pH 7.5). The column was incubated overnight at 4°C and unbound proteins were removed by washing with 10 column volumes of wash buffer (1× PBS, 1% Triton X-100, 2 mM DTT, 40 mM imidazole). The capsid protein was eluted using elution buffer (1× PBS, 1% Triton X-100, 2 mM DTT, 250 mM imidazole). The eluted fractions were analysed by 12% SDS-PAGE and those with the highest purity were pooled and dialyzed using a 3kDa cut-off membrane and used for western blotting. Blots were probed with anti-His antibody. For SARS-CoV2 N protein, the confirmed pET-28a-N construct was transformed into *E. coli* BL21DE3 cells and cultured in 500ml LB medium with 30µg/ml kanamycin under similar conditions. After induction with 1mM IPTG for 4 hours, the N protein was purified using Ni-NTA affinity chromatography with Tris buffer (pH 8.0).

### *In Vitro* pull-down assay

Ni-NTA beads were equilibrated with extraction buffer (PBS with 0.1% Triton X-100). HEK293 cell lysates were pre-cleared by incubating with equilibrated beads for 4 hours at 4°C. Ni-NTA beads pre-bound with purified c-anc protein were incubated with the pre-cleared lysate overnight at 4°C to allow interaction with host proteins. Beads were then washed three times with extraction buffer with 20 mM imidazole by centrifugation at 1,250L×Lg for 2Lmin. Bound proteins were eluted using a stepwise imidazole gradient (100-500LmM) in elution buffer. Eluted proteins were resolved by 12% SDS-PAGE and visualized with Coomassie Brilliant Blue staining. Protein bands of interest were excised and analysed by MALDI-TOF mass spectrometry (Galaxy International/Sandor Lifescience Pvt. Ltd. Hyderabad, India).

### Measuring mitochondrial membrane potential and ROS using live-cell imaging

HEK293 cells (1×10L) were seeded on 35mm glass-bottom dishes and transfected with plasmids encoding capsid variants. To measure the mitochondrial membrane potential, cells were incubated with 100nM tetramethyl rhodamine methyl ester (TMRM; Invitrogen, Cat. No. M20036) for 30Lmin at 37°C, washed with PBS and imaged live using a confocal microscope equipped with a 63× oil-immersion objective. To measure the reactive oxygen species (ROS) the cells were incubated with 5μM Cell ROX™ Deep Red reagent (Invitrogen, Cat. No. C10422) for 30Lmin at 37°C, rinsed with PBS and imaged live under identical acquisition settings. In drug treatment under infected conditions, Vero cells were pre-treated with 15μM UNA for 6Lh prior to infection and were infected with dengue virus infected cell supernatant. At 72Lh post-infection, cells were stained with TMRM or Cell ROX™ Deep Red as described above and imaged live.

### Mitochondrial stress assay

HEK293 cells were transfected with the recombinant constructs and Approximately 5000 cells were seeded onto poly-D-lysine (0.1% PDL)-coated XF mini 8-well plates and incubated for 48 hours at 37°C with 5% COL. Prior to assay, the cells were washed and incubated in Seahorse XF Assay Medium (Agilent) for 1 hour at 37°C in a non-COL incubator. OCR data was collected and analysed using Agilent Wave software v2.6.1.

Approximately 1 × 10L cells Vero cells (dengue infected and 15μM UNA treated) per well were seeded onto Seahorse XF mini cell culture microplates and were incubated for 72 hours. The mitochondrial stress test was conducted by sequential injections of mitochondrial inhibitors and uncouplers: 1μM oligomycin (ATP synthase inhibitor), 1μM FCCP (uncoupler) and a combination of 1μM rotenone and 1μM antimycin A (Complex I and III inhibitors). This sequence allowed us to determine the basal respiration, ATP production, maximal respiration, spare respiratory capacity and non-mitochondrial respiration.

### *In silico* based drug identification

As the crystal structure of the DENV1 c-anc protein was not available, we generated a homology model using SWISS-MODEL server[34]. Target-template alignment was performed using the ProMod3 tool available within the Swiss Model server. The model was built using PDB template 6VSO. The overall and per-residue quality of the models was evaluated using the QMEAN scoring function. Additionally, the validation of the c-anc protein structure was done using PROCHECK to validate the structural geometry using a Ramachandran Plot[35]. For docking analysis, the c-anc protein was prepared in AutoDock tools by removing water molecules, adding polar hydrogens, assigning Kollman charges and saving in PDBQT format[36]. Grid boxes were defined to encompass ligand-binding sites, maintaining default spacing. Virtual screening was performed using AutoDock Vina against the natural compound library available in the ZINC database[37], [38]. We choose natural compounds due to their structural diversity and well-established potential in antiviral drug discovery[39]. For this purpose, we retrieved natural compounds from the ZINC database and converted them into PDBQT format. Based on the binding energy, we selected the top 10 compounds and evaluated them for Lipinski’s rule of five using SwissADME[40]. Finally, the above top ten molecules were subjected to the Deep-PK web server to evaluate the pharmacokinetic properties, such as absorption, distribution, metabolism and excretion[41]. Complexes were visualized in PyMOL and analyzed in LigPlot+ to identify residues involved in protein-ligand interactions[42], [43]. The Ursonic acid (UNA) was selected in the above exercise.

The capsid protein-UNA complexes were processed using the Schrödinger Protein Preparation Wizard to ensure structural integrity and simulation readiness[44]. Key steps included the addition of missing side chains and loops, optimization of hydrogen-bonding networks, correction of protonation states at pH 7.4 and removal of non-essential water molecules. The prepared complexes were embedded into a rhombic dodecahedral box using the System Builder tool in Schrödinger. The OPLS4 force field was applied and the systems were solvated using TIP4P water models. Physiological ionic conditions were replicated by adding 0.15M NaCl and appropriate counter-ions to neutralize the system. Each complex underwent a 100 nanosecond MD simulation using the Desmond module. Snapshots were saved every 2ps, resulting in ∼10,000 frames per simulation. Root Mean Square Deviation (RMSD) analyses were performed using Maestro’s Simulation Interaction Diagram (SID) tool. The initial frame of the trajectory was used as the reference and protein backbone atoms were monitored to evaluate global conformational drift. In parallel, ligand RMSD was computed to evaluate the positional retention of UNA within the binding site. We have carried out a similar analysis using the SARS-CoV-2 nucleocapsid protein and UNA.

### Intrinsic fluorescence measurements

The intrinsic fluorescence emission spectra of the capsid proteins were recorded using a Jasco Spectrofluorometer (FP-8500) with a protein concentration (c-anc of DENV1 and N of SARS CoV2, pH 7.4) of 10μM in 1× PBS. Measurements were performed in a 10 mm path length cuvette at 25°C. Protein excitation was carried out at 280 nm and emission spectra were recorded. The scanning speed was set to 100 nm/min, with a slit width of 5 nm. For UNA (Cat# CS-5796, ChemScene) binding studies, purified capsid proteins were incubated with varying concentrations of UNA.

### Cell morphology analysis

Vero cells seeded in a 12 well plate were pre-treated with 5, 10 and 15µM concentrations of UNA, followed by dengue infection. After 72 hours, the plates were collected and images were captured under an inverted bright field microscope (LYNXR^®^ Lawrence & Mayo) at 10X.

### DENV-1 infection and UNA treatment in mice

Dengue virus infection was established as previously described[45], [46], [47]. Female C57BL/6 mice (Hylasco Biotechnology Pvt.Ltd, India) were fed with a high-fat diet (VRK nutritional solutions, Hyderabad, India) for 4 weeks prior to infection. The diets were maintained throughout the study. After feeding, the mice were administered 1mg of anti-IFNAR antibodies by intraperitoneal route. 24 hours later, the mice were infected with 100 µl DENV1 (2 x 10^4^ FFU) by intradermal and 50µl through subcutaneous at the plantar region of the hind paw. The animals were randomized into three groups, each comprising four mice: mock-infected, virus-infected and virus-infected and treated with 100 mg/kg UNA. For the haematological study, blood samples were collected from the retro-orbitals of mice under anaesthesia on 3, 5 and 7 days post infection. In an 0.5 ml Eppendorf tube, 300µl of blood sample was mixed with 20µl of freshly prepared 5% dipotassium ethylenediaminetetraacetic acid (K2EDTA), an anticoagulant. For viral RNA analysis, approximately 150 µl serum was separated from the remaining blood samples, aliquoted and stored at -80°C.

### RT PCR of viral RNA

Viral RNA was isolated from both infected cell supernatants and serum samples collected from infected mice. cDNA synthesis was performed as described earlier. For RT-PCR, dengue virus 5’ or 3’UTR primers and GCC biotech master mix were used. The amplification mixture consisted of 4 µl of cDNA, 5 µl of master mix and 200nM each of forward (CAATATGCTGAAACGCGAGAGAAA) and reverse (CCCCATCTATTCAGAATCCCTGCT) 5’UTR primers and forward (GGTAAGCCAACACACTCATGAAA) and reverse primers (TGCCTGGAATGATGCTGTAGA) of 3’UTR were used. The mixture was subjected to 95°C for 5 minutes, followed by 35 cycles of 95°C for 1 minute, 58°C for 30 seconds (for 3’UTR 56°C for 50 seconds) and 72°C for 30 seconds (for 3’UTR 72°C for 1 minutes), with a final extension at 72°C for 2 minutes (for 3’UTR 72°C for 5 minutes). The amplified products were analysed on 1% agarose gel electrophoresis.

### Haematological and histopathological assays

Parameters such as white blood cells, neutrophils, lymphocytes, mixed cell, red blood cells, hemoglobin, hematocrit, mean corpuscular hemoglobin, mean corpuscular hemoglobin concentration, RDW-CV, RDW-SD, platelet count, mean platelet volume and plateletcrit were analysed using an automated haematology analyzer (BHA-3000 VET Automatic Hematology Analyzer). For histological analysis, spleen tissues were dissected, cleaned to remove excess fat and were fixed in 2% paraformaldehyde at room temperature for 1 hour. Then rinsed under running tap water for 5 minutes and dehydrated through a graded ethanol series (70%, 80%, 95%; 5 minutes each) followed by three changes of 100% ethanol (5 minutes each). The tissues were cleared in xylene twice for 5 minutes each and infiltrated with molten paraffin three times for 5 minutes each before embedding in paraffin blocks. Paraffin-embedded tissues were sectioned at 5-8Lμm using a microtome and floated on a 40°C water bath containing distilled water. Sections were mounted on glass slides and slides were deparaffinized in xylene (2 × 5 minutes). The sections were rehydrated through a graded ethanol series (100% ×2, 95%, 70%, and 50%; 3 minutes each), and washed in PBS (2 × 5 minutes). Endogenous peroxidase activity was quenched by incubating sections in 3% HLOL in methanol at room temperature for 10 minutes, followed by PBS washes (2 × 5 minutes). Antigen retrieval was performed using 10LmM citrate buffer (pH 6.0). Then the slides were placed in a staining container with buffer and heated to 95-100°C for 10 minutes. Slides were then cooled at room temperature for 20 minutes and rinsed in PBS (2 × 5 minutes). To reduce nonspecific binding, sections were blocked with 0.5 % BSA in PBS for 30 minutes. Then the section were incubated in Interferon Gama protein 1 antibodies (IFNGR1) antibodies for 1 hour at room temperature in a humidified chamber. Following PBS washes (2 × 5 minutes), sections were incubated with HRP conjugated secondary antibody for 30 minutes at room temperature followed by PBS washes (2 × 5 minutes). Color development was performed using freshly prepared DAB substrate solution. Slides were washed in PBS (3 × 2 minutes). For counterstaining, slides were immersed in hematoxylin for 1–2 minutes, rinsed under running tap water for 10 minutes, dehydrated through a graded ethanol series (95% ×2, 100% ×2; 5 minutes each) and cleared in xylene (3 × 5 minutes). Finally, the stained sections were visualized under a light microscope.

### C-anc-SNTA1 docking

The amino acid sequences of [DENV1 (ASD49619.1), DENV2 (AYE66919.1), DENV3 (YP_001621843.1), DENV4 (ARB18129.1), ZIKV (YP_002790881.1), YFV (NP_041726.1), JEV (NP_059434.1) and HCV (NP_671491.1), CHIKV (NP_690589.2), ASFV (NP_042775.1) and SARS-CoV2 (YP_009724397.2)] were retrieved from the NCBI protein database. These sequences were used to obtain corresponding capsid or nucleocapsid protein structures either directly from the Protein Data Bank (PDB) or by homology modeling using the SWISS-MODEL server. All modelled structures were validated using PROCHECK based on Ramachandran plot analysis. We used ClusPro 2.0 web server to assess the binding of SNTA1 with capsid proteins[48]. Protein–protein interactions were visualized using PyMOL.

### UNA interactions with capsids of other viruses

Capsid protein structures of flavi (DENV1, ZIKV, JEV, YFV and HCV) and non-flaviviruses (CHIKV, SARS-CoV2 and ASFV) were sourced from the PDB or modelled via SWISS-MODEL server then validated using PROCHECK through Ramachandran plots. The structure-UNA docking was carried as described earlier.

## Results

### Dengue virus capsid protein exists in multiple forms

The whole-cell lysates of DENV1 infected K562 cells were collected at 1, 6, 12, 24, 48 and 72 hours post-infection and analysed by western blot using the anti-capsid antibodies. The data revealed the presence of multiple bands with the molecular weights of approximately 12, 21, 25 and 32 kDa, suggesting that they are time dependent processed forms of the polyprotein (Figure 1A [i]). The molecular weights indicate that the multiple forms could represent the capsid (12 kDa), c-anc (21 kDa), c-anc-pr (25 kDa), c-anc-pr (32kDa). The mature capsid band (∼12kDa) was consistently observed from 1 to 72 hours and the c-anc (∼21kDa) appeared after 12 hours and remained detectable up to 72 hours. The c-anc-pr (∼25kDa) and c-anc-prM (∼32 kDa) forms was detected specifically at 72 hours. The quantification of these bands at different time points supported the above pattern of proposed capsid processing (Figure 1A [iii]). Based on these observations and the reported cleavage sites, we hypothesized the possible cleavage intermediates of capsid protein and developed the recombinant constructs as described in the methods for further analyses (Figure 1B; Figure S1A).

**Figure 1.**
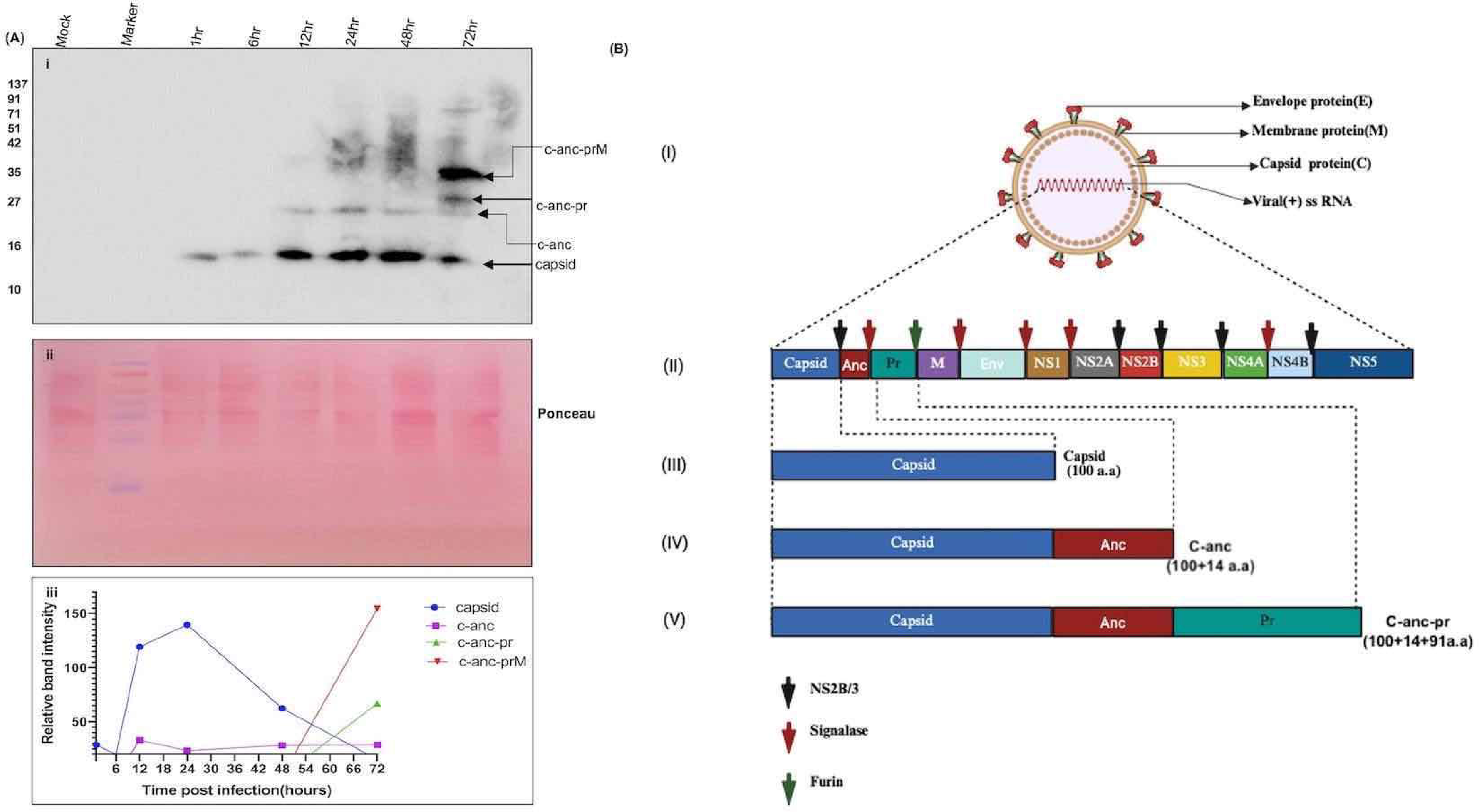
Dengue virus capsid protein processing during infection in K562 cells. (A) (i) DENV1-infected K562 cells were harvested at the indicated time points post-infection and probed with anti-capsid antibody; (ii) Ponceau S staining of the same membrane (iii) Intensity profiles for the expression levels of capsid, c-anc, c-anc-pr and c-anc-prM. (B) (i) Structure of the dengue virus; (ii) Graphical representation (created using BioRender tool) of the dengue virus polyprotein, illustrating the structural proteins (capsid, pre-membrane and envelope) and non-structural proteins (NS1-NS5); based on possible forms observed in figure A, we speculate the existence of three processed forms of capsid protein; (iii) Mature capsid protein (100 amino acids); (iv) c-anc protein (114 amino acids); and (v) c-anc-pr protein (206 amino acids). Arrows on the polypeptide indicates the cleavage sites.

### Dengue c-anc and c-anc-pr proteins localise to the mitochondria

There is literature evidence that virus proteins that exist in multiple forms show the tendency to localize into different sub-cellular organelles and influence their homeostasis[8], [9], [21], [22], [49]. Hence, we aimed to investigate the mitochondrial and nuclear localization of different capsid forms observed in this study. In this direction, HEK293 cells were transfected with GFP-tagged constructs expressing the c, c-anc and c-anc-pr. Confocal microscopy of transfected cells demonstrated the mitochondrial localization for c-anc, c-anc- pr and nuclear localization for capsid protein (Figures 2A [i-xii]; S1B and S1C). The nuclear localization of the capsid was already reported [50]. The c-anc showed the strongest localization to the mitochondria, while c-anc-pr also localized to mitochondria but with comparatively weaker signals. These observations were further supported by Pearson correlation analysis data developed from the above experiment, where the c-anc and c-anc-pr forms exhibited high correlation coefficients, indicating strong mitochondrial localization. In contrast, the c form showed a lower correlation coefficient, suggesting reduced/no mitochondrial association (Figure 2B [v]). Multiple sequence alignment of capsid proteins from DENV1-4 revealed the conservedness among the serotypes, hence the existence of multiple capsid forms and their sub-cellular localizations in the case of other serotypes could also be similar to DENV1. Since the c-anc and c-anc-pr showed similar subcellular localizations, we continued with c-anc only for further analyses.

**Figure 2.**
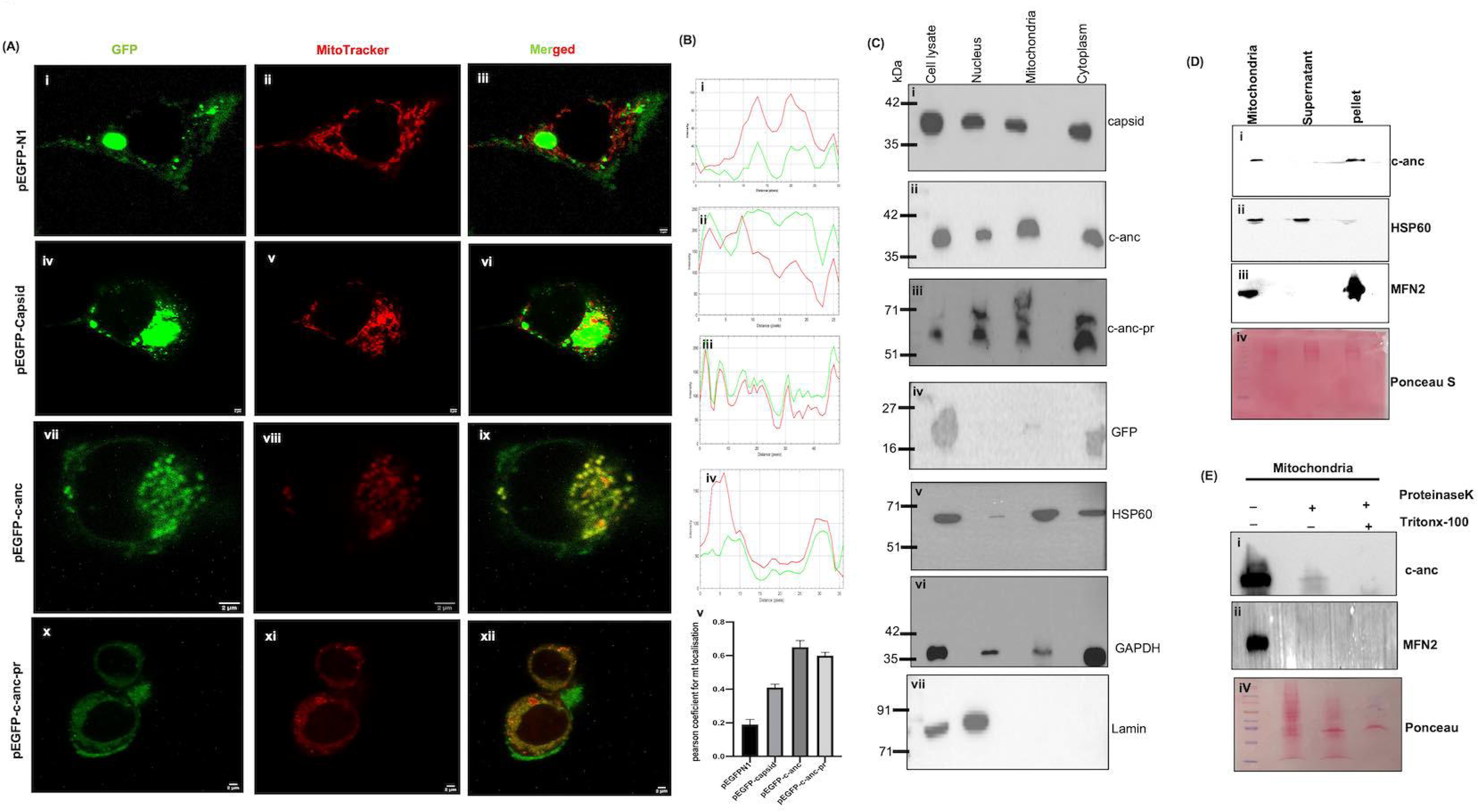
c-anc protein localises to mitochondria. (A) Confocal images acquired at 63× magnification (scale bar: 2Lµm) show the localization of (i-iii) pEGFP-N1 vector; (iv-vi) pEGFP-c; (vii-ix) pEGFP-c-anc and (x–xii) pEGFP-c-anc-pr, along with the mitochondrial localization marker, i.e., MitoTracker® Red CMXRos. (B) (i–iv) Represents the fluorescence intensity profiles of GFP and mitochondria; (v) Graph representing the Pearson correlation coefficient of GFP, c, c-anc and c-anc-pr with mitochondria. (C) (i–iv) Subcellular fractions from HEK293 cells transfected with pEGFP-c, pEGFP-c-anc, and pEGFP-c-anc-pr constructs and pEGFPN1 vector and probed with anti-GFP antibody;(v) reprobed with anti-HSP60 antibody; (vi) anti-GAPDH antibody; (vii) anti-Lamin antibody. (D) (i–iii) Alkali-treated mitochondrial fractions from pEGFPN1-c-anc transfected HEK293 cells, probed with anti-GFP, anti-HSP60 and anti-MFN2 antibodies; (iv) Ponceau S stained membrane, probed and reprobed with the above antibodies. (E) (i–iii) Proteinase K treated Mitochondrial fractions from pEGFP-c-anc transfected cells were immunoblotted using anti-GFP and anti-MFN2; (iv) Ponceau S stained membrane, probed and reprobed with the above antibodies.

In order to confirm the above data, subcellular fractionations were performed. The mature capsid protein was present in all fractions, with predominant enrichment in the nuclear extract and minor presence in the mitochondrial fraction. The c-anc and c-anc-pr forms were also detected across all fractions but showed higher abundance in the mitochondrial fraction, indicating mitochondrial localization (Figure 2C [i-iv]). Among the fractionation markers, HSP60 was predominantly enriched in the mitochondria, Lamin A/C was confined to the whole cell and nuclear extracts and GAPDH was detected in all fractions, with highest levels in the cytoplasmic extract, confirming the authenticity of fractionation (Figure 2C [v-vii]). Further, localization findings were validated in dengue virus-infected Vero cells, data suggested that both fluorescence microscopy and immunoblotting confirmed mitochondrial localization of the endogenous c-anc protein (Figure S2A [i-viii]; S2B [i-iii]). To determine the sub-mitochondrial localization of the c-anc, an alkali extraction assay using sodium carbonate (NaLCOL) was performed. Immunoblotting of the samples from the above experiment revealed that c-anc is predominantly enriched in the pellet fraction, similar to membrane-integrated proteins (MFN2), suggesting its localization to the mitochondrial membrane (Figures 2D [i-iv]).

To further evaluate its association with outer or inner membranes of mitochondria, we isolated mitochondria from c-anc transfected cells and treated with proteinase K (Pro K). We observed a reduction in band intensity due to Pro K digestion, suggesting its localization to the outer membrane. As expected, MFN2 (outer mitochondrial membrane) protein was also found to be digested by Pro K, confirming the accessibility of the outer membrane (Figure 2E [i-iii]).

### c-anc protein triggers changes in mitochondrial homeostasis

Confocal microscopy observations revealed that expression of c-anc and c-anc-pr induced mitochondrial fragmentation in HEK293 cells, while the mitochondria in control cells remained elongated and tubular, a characteristic feature of regular mitochondria (Figures 3A [i -viii]). Among all constructs, c-anc expression caused the most significant morphological disruption. Transmission electron microscopy (TEM) studies confirmed these findings as c-anc transfected HEK293 cells exhibited fragmented mitochondria, which were further strengthened by the similar effects observed in Vero cells infected with DENV1 (Figures 3B [i-viii]).

**Figure 3.**
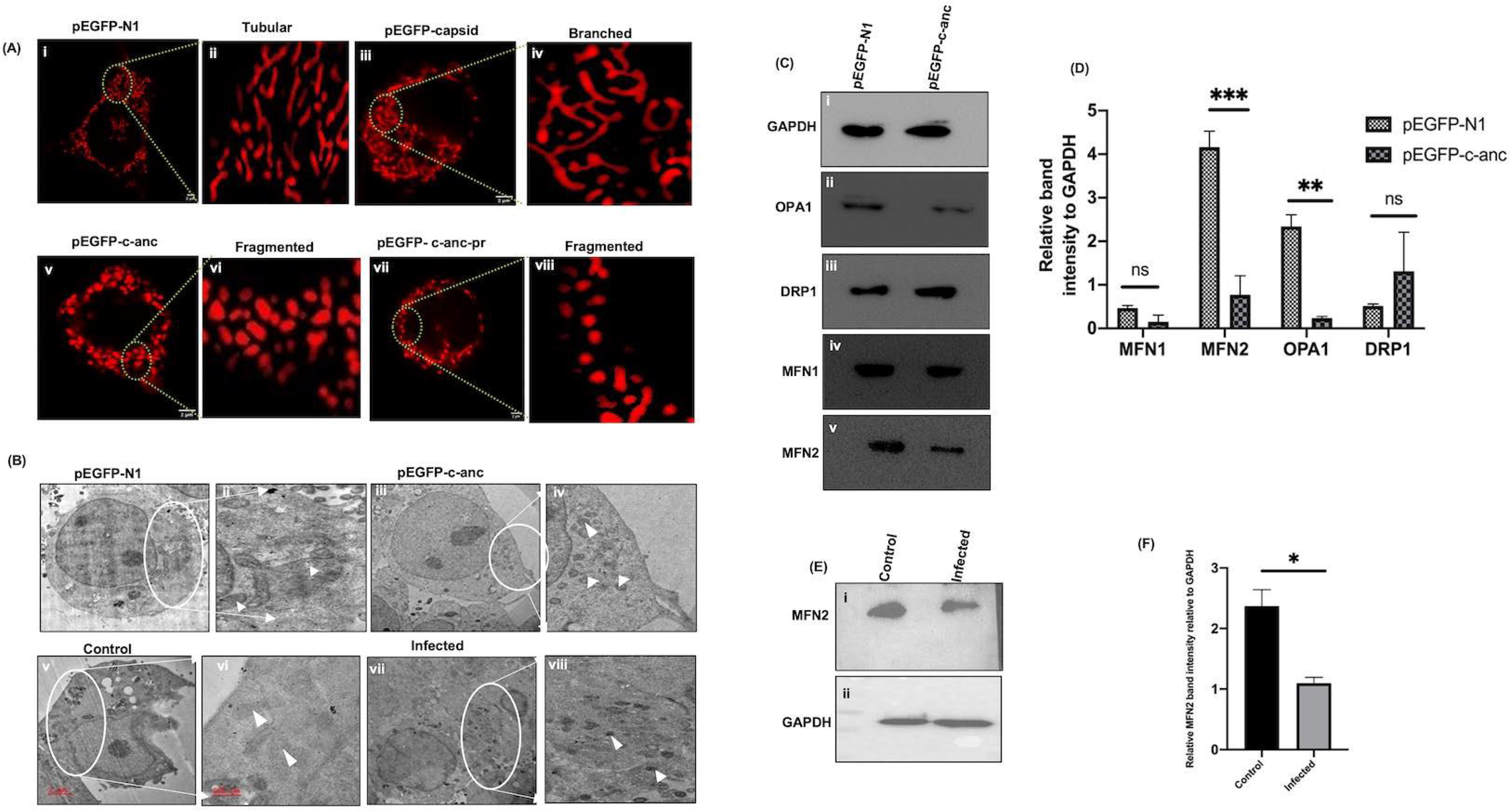
Dengue virus capsid protein variants trigger changes in mitochondrial homeostasis. (A) Confocal images of HEK293 cells transfected with pEGFPN1 and capsid variants, stained with TMRM dye; Mitochondria in (i–ii) pEGFP-N1 vector ; (iii-iv) capsid; (v–vi) c-anc; and (vii-viii) c-anc-pr transfected cells, along with their respective enlarged views. Scale bar = 2Lµm. (B) Transmission electron microscopy (TEM) of transfected HEK293 and infected Vero cells; (i–ii) pEGFP-N1 vector; (iii-iv) pEGFP-c-anc; (v–vi) Mock; and (vii-viii) Infected Vero cells, along with their respective enlarged views. Scale bar = 2Lµm (C) (i–v) Western blot analysis of HEK293 cells transfected with pEGFP-N1 and pEGFP-c-anc, using antibodies against GAPDH, OPA1, DRP1, MFN1 and MFN2. (D) The bar graph shows the relative band intensities of OPA1, DRP1, MFN1 and MFN2 normalized to GAPDH. (E) Western blot analysis of control and virus-infected Vero cells at 72 hours post-infection, probed with antibodies against (i) MFN2 and (ii) GAPDH. (F) Bar graph showing the mean MFN2 expression levels normalized to GAPDH in control and infected cells.

To investigate these alterations further at the protein level, we analysed the expression of key mitochondrial fusion/fission proteins by western blotting. In this direction, the levels of mitochondrial fusion proteins OPA1 and MFN2 were found to be significantly reduced (Figures 3C [i-v] and 3D). Similar patterns were observed in DENV1-infected Vero cells, where MFN2 expression was found to be reduced (Figures 3E [i and ii] and 3F). The change in DRP1 levels was insignificant in c-anc expressing cells, suggesting that mitochondrial fission was not actively involved.

Further, RNA-seq analysis of cells transfected with c-anc revealed a significant change in gene expression. In total, 3,674 genes were found to be differentially expressed, of which 1,872 genes were found to be upregulated and 1,802 were downregulated. A heatmap of the top 50 differentially expressed genes showed distinct expression patterns between the two groups (Figure S3A). The volcano plot further highlighted these differences, showing statistical significance for each gene (Figure S3B). To validate the transcriptomic findings, real-time quantitative PCR (RT-qPCR) was performed for a subset of genes from the above top 50 that are involved in mitochondrial function. Consistent with the RNA-seq data, ATP6, CO3, ND2 and CYB were found to be affected (Figure S3C). The literature indicate that the above affected genes involve in the metabolism of the mitochondria[33].

### c-anc interacts with SNTA1

The literature suggest that dengue virus capsid protein interacts with nucleolin and localizes to the nucleus[20], [50]. In order to identify any host factors that interact with the c-anc protein, we performed a pull-down assay. A protein band with a molecular weight between 50 and 70 kDa was observed in the eluates, excised and analyzed by mass spectrometry (MS-TOF) (Figure S4A [i]). Alpha-1 syntrophin (SNTA1) showed the second highest score (34) among the identified and functionally annotated proteins (Figure S4C). Although an uncharacterized protein had a slightly higher score but information regarding that protein was not available. The western blot analysis of the above eluates using anti-SNTA1 antibody confirmed the above MS-TOF result (Figures 4A [i-iii]). In order to confirm the c-anc-SNTA interactions, we conducted *in silico* protein–protein docking analysis using the ClusPro server. The docking results revealed a favourable and stable interaction between c-anc and SNTA1, with a minimum binding energy of −1144.8 kcal/mol (Table S2).

**Figure 4.**
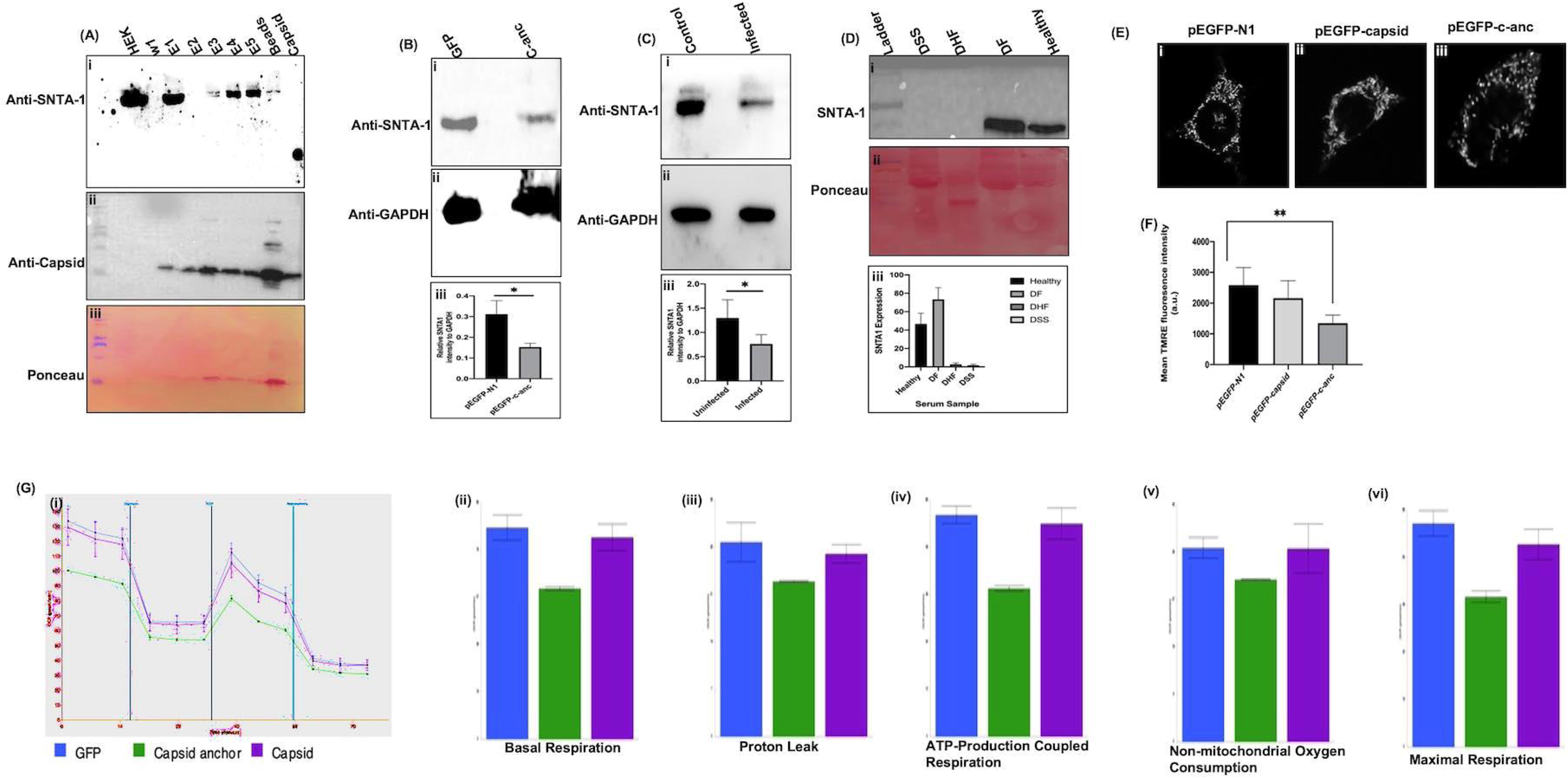
Dengue virus c-anc interacts with host SNTA1 protein and downregulates its expression in transfected, infected cells and clinical samples and impairs mitochondrial bioenergetics. (A) (i) Western blot analysis of pull-down assay using purified c-anc and HEK cell lysates showing SNTA1 in the elutions; (ii) Anti-capsid antibody confirms the presence of c-anc in the pull-down fraction; (iii) Ponceau S stained membrane probed and reprobed with the above antibodies. (B) Western blot analysis of c-anc transfected cells; (i) Blot probed with anti-SNTA1; (ii) Same blot stripped and reprobed with anti-GAPDH as a loading control; (iii) Bar graph showing SNTA1 expression levels normalized to GAPDH. (C) (i) Western blot analysis using anti-SNTA1 antibody in infected cells; (ii) Blot probed with anti-GAPDH; (iii) Bar graph showing SNTA1 expression levels normalized to GAPDH. (D) Western blot of serum samples from healthy and dengue infected patients (DF, DHF and DSS); (i) anti-SNTA1; (ii) ponceau stained membrane; (iii) Bar diagram represents the relative densities of SNTA1 proteins in clinical samples. (E) Representative images for mitochondrial membrane potential in TMRM stained HEK293 cells (i-iii). Cells were analysed by confocal microscopy. (F) Quantification of TMRM fluorescence intensity of the above confocal images. (G) (i) Agilent Seahorse XFp Mitochondrial Stress Test showing mitochondrial bioenergetics in cells transfected with pEGFP-N1vector, pEGFP-capsid and pEGFP-c-anc represented by oxygen consumption rate (OCR) over time during sequential injection of three drugs: Oligomycin, FCCP and Rotenone/Antimycin; Graphs showing oxygen consumption rate (OCR) over time (ii) Basal respiration; (iii) Proton leak; (iv) ATP-production coupled respiration; (v) Non-mitochondrial oxygen consumption; (vi) Maximal respiration.

### c-anc affects SNTA1 levels in transfected, infected cells and patient serum samples

The western blot analysis of c-anc transfected HEK293 and DENV1-infected Vero cells showed reduced levels of SNTA (Figures 4B [i-iii]; 4C [i-iii]). Further, SNTA1 protein levels in clinical samples were found to be significantly reduced or absent in serum samples from individuals with dengue hemorrhagic fever (DHF) and dengue shock syndrome (DSS) (Figures 4D [i-iii]). Total fifteen clinical samples with DF, DHF and DSS were included for analysis. The docking studies indicate that SNTA interacts with the c-anc of other serotypes of dengue virus as well (Table S2).

### c-anc impairs mitochondrial bioenergetics

We also performed mitochondrial membrane potential (ΔΨm) assay using the ΔΨ-dependent fluorescent dye TMRM. Real-time confocal microscopy revealed a reduction in mitochondrial membrane potential in cells expressing the c and c-anc constructs (Figure 4E [i-iii] and 4F). Then, we assessed capsid and its variant’s impact on mitochondrial function using the Seahorse XF Mito Stress Test in HEK293 cells. Five different key respiratory parameters were evaluated to determine bioenergetic performance. Basal respiration and Proton leak were found to be reduced in c-anc expressing cells compared to both vector and capsid expressing cells. ATP production was also found to decline in c-anc transfected cells upon addition of oligomycin, an ATP synthase inhibitor. Maximal respiration was calculated following uncoupler FCCP treatment, c-anc-expressing cells were found to fail to exhibit the expected increase in OCR, indicating the compromised electron transport chain (ETC) integrity and limited substrate availability. Similarly, non-mitochondrial oxygen consumption, measured after inhibition of complex III by antimycin and rotenone, was decreased in c-anc-transfected cells (Figure 4G [i-vi]).

### Ursonic acid interacts with c-anc protein *in silico*, *in vivo* and *ex vivo*

The 3D structure of the DENV1 c-anc protein was successfully modelled using the SWISS-MODEL server (Figure 5A). The Ramachandran plot analysis revealed that 99.2% of residues were in the most favourable, 0.8% in the additionally allowed regions and none in the generously allowed or disallowed regions (Figure 5B). We docked compounds from the ZINC database with the DENV1 capsid protein and selected the top 10 molecules based on Lipinski’s Rule of Five, ADMET properties, and binding energy values. Among these, UNA stood out topmost due to its strong binding and fulfilling all parameters of the Lipinski rule (Table 1 and Table S1). Molecular dynamics simulations suggest that UNA exhibits stable and consistent interactions with the c-anc protein. The Root Mean Square Deviation (RMSD) c-anc-UNA showed stabilized complex around 6.8 Å, indicating a stable conformation with minimal structural deviation throughout the simulation (Figure 5D). The intrinsic fluorescence of the purified c-anc protein showed a dose-dependent decrease in intensity upon interaction with increasing concentrations of UNA (Figure 5E and 5F). A reduction in fluorescence was observed starting at concentrations of 2nM UNA, with a more pronounced effect at higher concentrations indicating a dose dependent interaction between UNA and the c-anc protein (Figure 5F).

**Figure 5.**
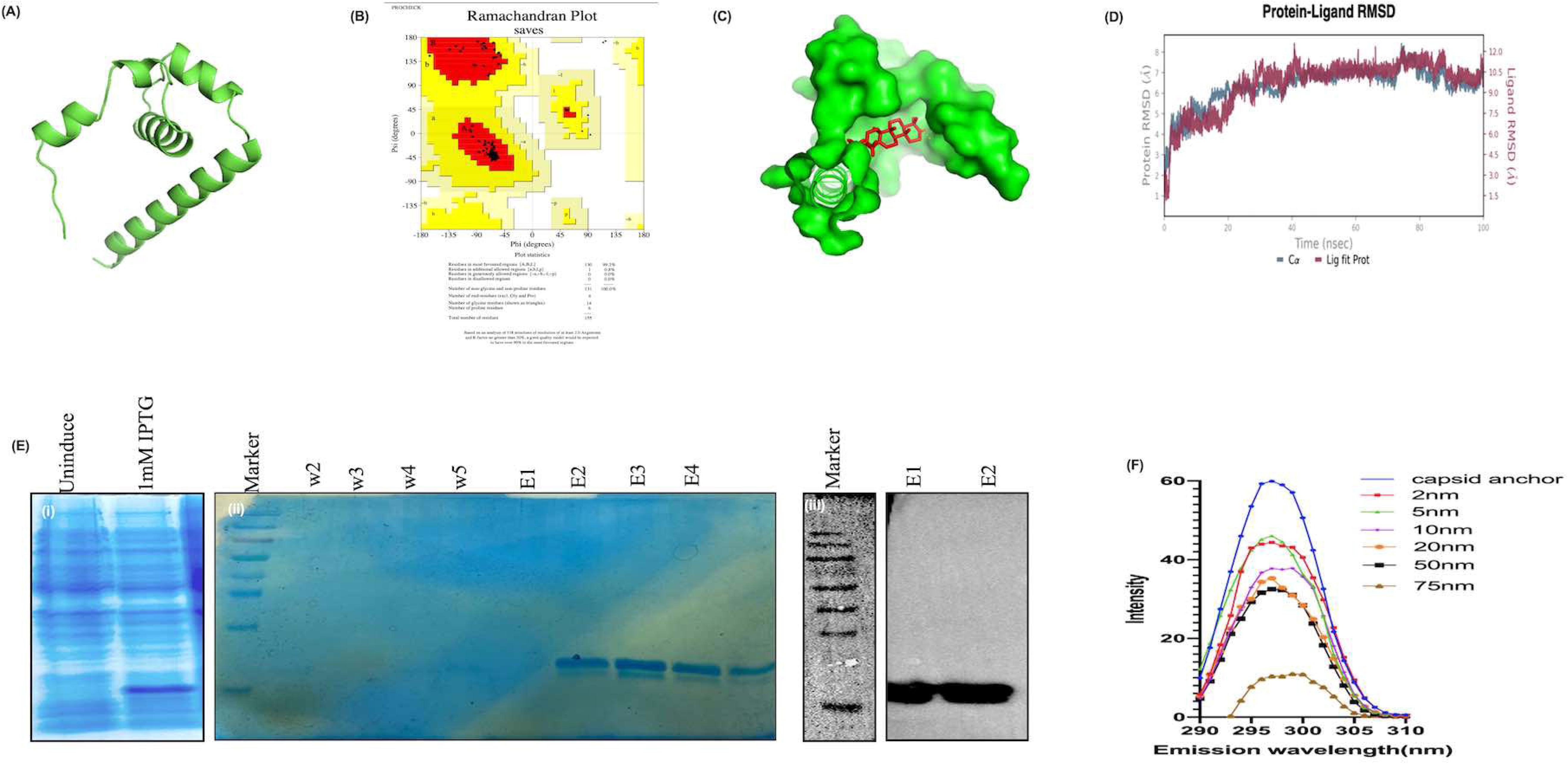
Ursonic acid interacts with purified c-anc protein. (A) 3D structure of DENV1 c-anc protein. (B) Ramachandran plot of modelled c-anc protein. (C) Molecular docking showing ligand UNA (stick model) binding to the capsid protein. (D) RMSD analysis of the UNA-bound c-anc illustrates the changes in protein-ligand during the molecular dynamics simulation of the c-anc. (E) Expression and purification of the DENV1c-anc Protein; (i) Lane 1-uninduced sample, Lane 2 expressed c-anc protein after induction; (ii) Lane 1-Pretein ladder, Lane 2 to 7 Elution fractions (iii) Western blot validation of c-anc expression in selected elution fractions. Lane 1: molecular weight marker; Lanes 2–3: purified protein fractions probed with anti-His antibody. (F) Effect of UNA on the intrinsic fluorescence of the c-anc protein in the presence and absence of UNA as indicated with different colours.

**Table 1:**
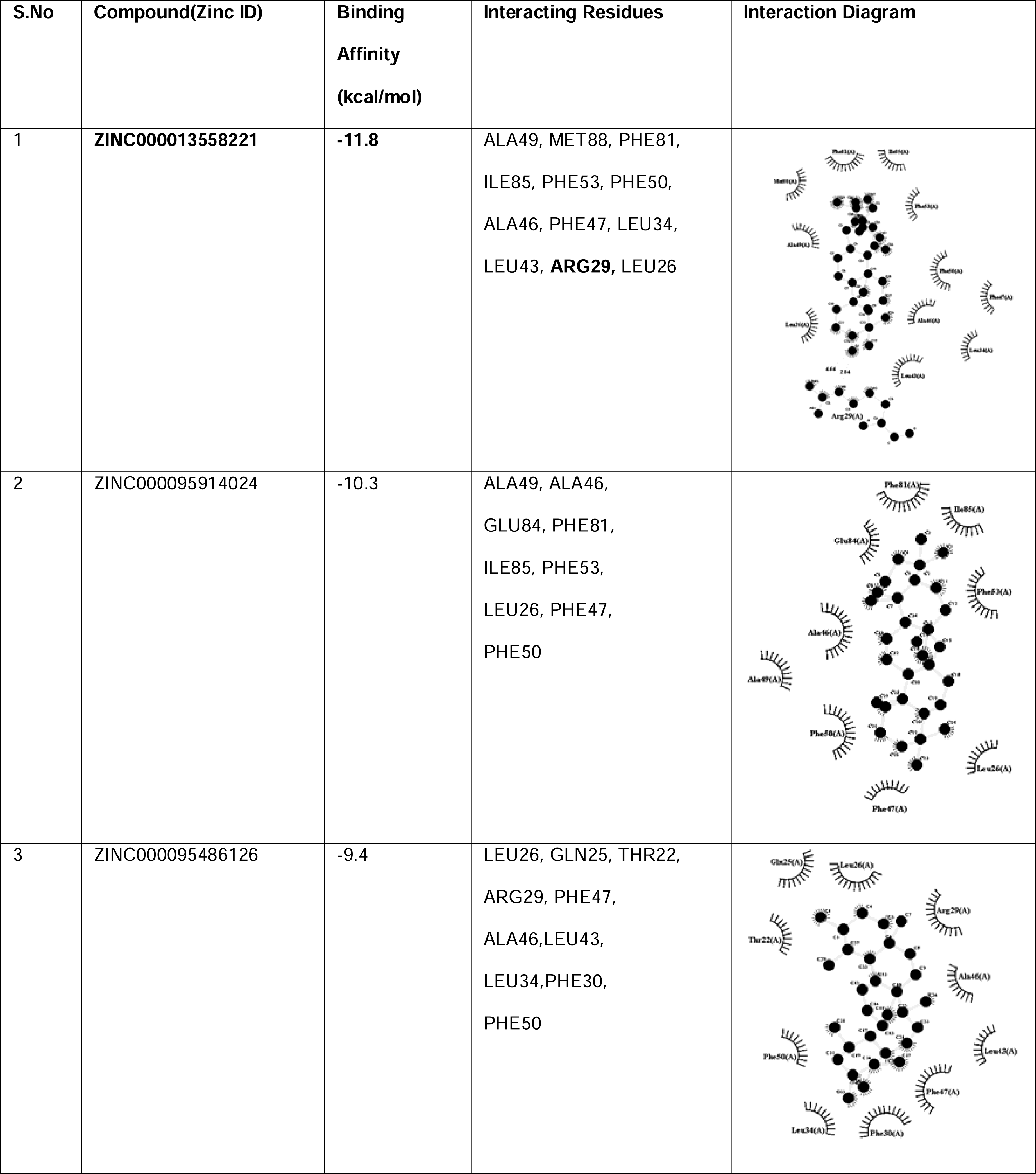
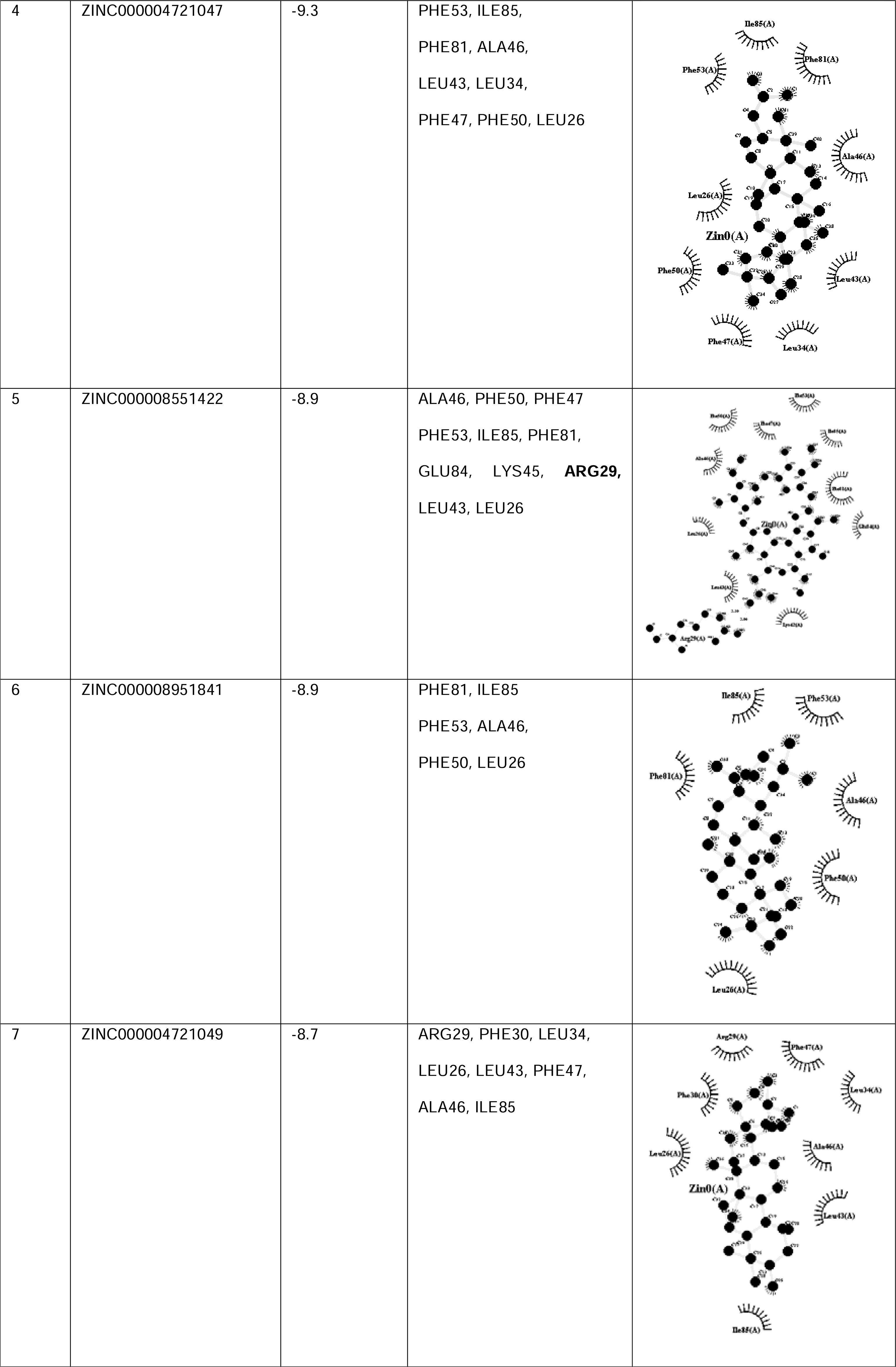
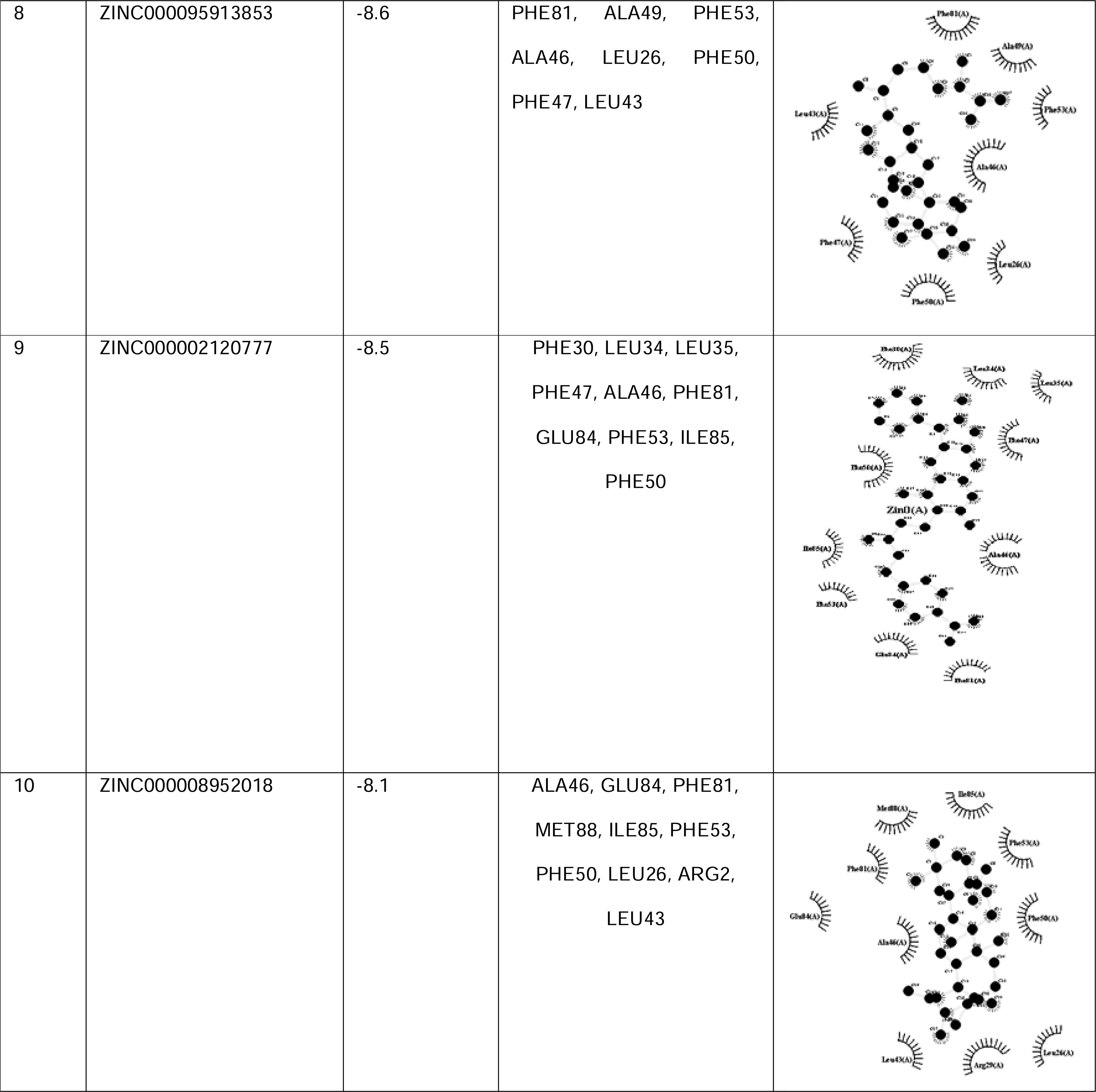
Protein-Ligand interaction analysis of the top 10 compounds with c-anc protein.

To confirm the inhibitory effect of UNA on DENV1 multiplication, Vero cells were treated with varying concentrations of UNA (0, 5, 10, and 15μM) for 6 hours, followed by infection with DENV1 supernatant. After 72 hours post-infection, confocal microscopy images revealed a dose-dependent reduction in fluorescence in dengue-positive cells with increasing concentrations of UNA (Figure 6A [i-xv]). At 15μM concentrations, UNA exhibited significant inhibition of dengue replication. The quantification of dengue related fluorescence in cells confirmed these findings (Figure 6B). To further assess the effect on viral replication, total RNA was extracted from the cell supernatants and analyzed by qPCR using DENV1 3’UTR-specific primers. The results indicated a significant reduction in viral load in the supernatants of cells treated with UNA (Figure S5A). Western blot analysis was performed to measure the expression levels one of the viral proteins (NS2BNS3) in Vero cells treated with UNA. The results revealed a dose-dependent reduction in NS2BNS3 protein levels with the highest inhibition at 15μM UNA (Figure 6C [i-iii] & D). Deformed cell morphology, rounding/clubbing of cells and the cytopathic effect were observed after the DENV1 infection, but were found to be reverted to normal conditions upon UNA treatment (Figure S5B [i-v]). To investigate the functional consequences of c-anc on mitochondrial activity in presence of UNA, we evaluated the mitochondrial membrane potential (ΔΨm) and ROS levels. Real-time microscopy revealed a reduction in mitochondrial membrane potential in cells transfected with c-anc. The drug treated c-anc transfected HEK293 cells demonstrated restoration of mitochondrial potential and ROS levels (Figure S5C and S5D). The above parameters showed a similar trend in dengue infected and drug treated Vero cells (Figure 6E [i-iii],6F and S5E). Further, pre-treatment with UNA restored mitochondrial functions (OCR across basal respiration, ATP production, non-mitochondrial respiration, spare respiratory capacity and maximal respiration) when compared to only virus infected conditions (Figure 6G [i-vi]). The docking studies indicate that UNA binds with the c-anc of other serotypes of dengue virus as well (Table S3).

**Figure 6.**
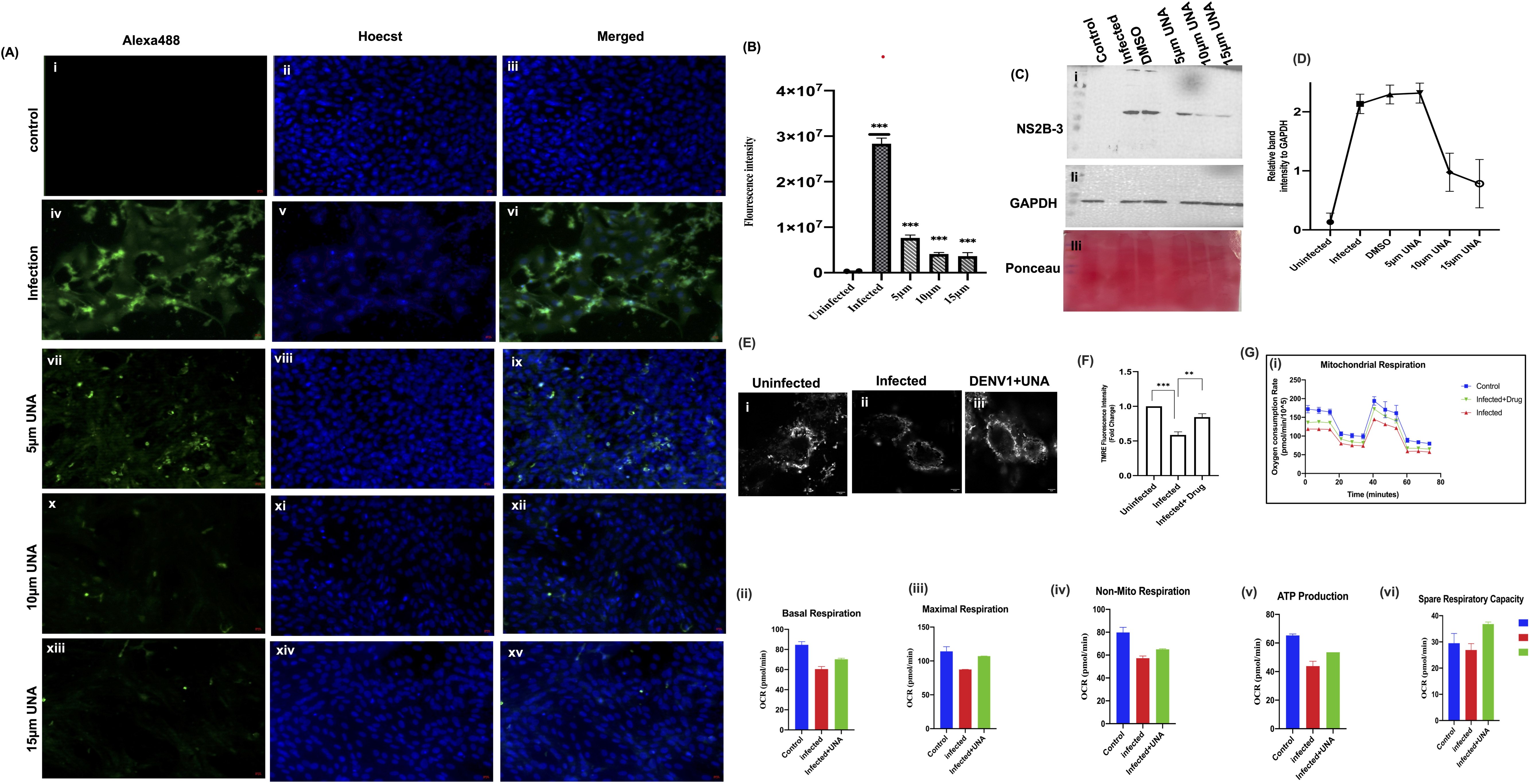
Ursonic acid reduces dengue virus multiplication and restores mitochondrial function in infected Vero cells. (A) Immunofluorescence images showing NS2BNS3 protein expression in Vero cells infected with DENV1 and pre-treated with UNA; (i–iii) Uninfected ; (iv–vi) Infected cells; (vii–ix) 5 µM ; (x-xii) 10 µM ; and (xiii–xv) 15 µM UNA-treated cells. (B) Bar graph representing total fluorescence intensities from the above experiment. (C) Western blot analysis of viral protein expression upon UNA treatment: (i) Immunoblot showing NS2BNS3 protein levels in Vero cells treated with UNA at 5,10 and 15LμM concentrations; (ii) anti-GAPDH antibody; (iii) Ponceau S membrane probed and reprobed with above antibodies. (D) Quantification of NS2BN3 band intensity normalized to GAPDH. (E) Representative images for mitochondrial membrane potential in Vero cells; (i-iii) TMRM fluorescence intensity in uninfected, DENV1-infected and UNA-treated infected cells (F) Bar graph representing the quantification of TMRM fluorescence intensity. (G) (i)Agilent Seahorse XFp Cell Mito Stress Test showing real-time oxygen consumption rate (OCR) in mock, DENV1-infected and ursonic acid-treated DENV1-infected Vero cells plotted over time. Bar graphs showing (ii)basal respiration (iii)maximal respiration (iv)non-mitochondrial respiration (v)ATP production and spare (vi)respiratory capacity.

### Ursonic acid inhibits dengue virus multiplication in mice

In a pilot experiment, carried out in DENV1 infected and UNA treated mice, the clinical parameters (body weight and temperature) were assessed (Figure S5F [i-ii]). Further, the hematological studies revealed a statistically significant improvement, particularly in the platelet count (Figure S5F [i-vi]). Immunohistochemistry (IHC) of spleen sections showed low levels of IFNGR1 protein in infected mice, while the mice treated with UNA showed much higher levels of IFNGR1 protein (Figure 7A and 7B). RT-PCR analysis showed a significant reduction in viral RNA levels in the blood sample of UNA-treated compared to untreated infected mice (Figure 7C [i and ii]).

**Figure 7:**
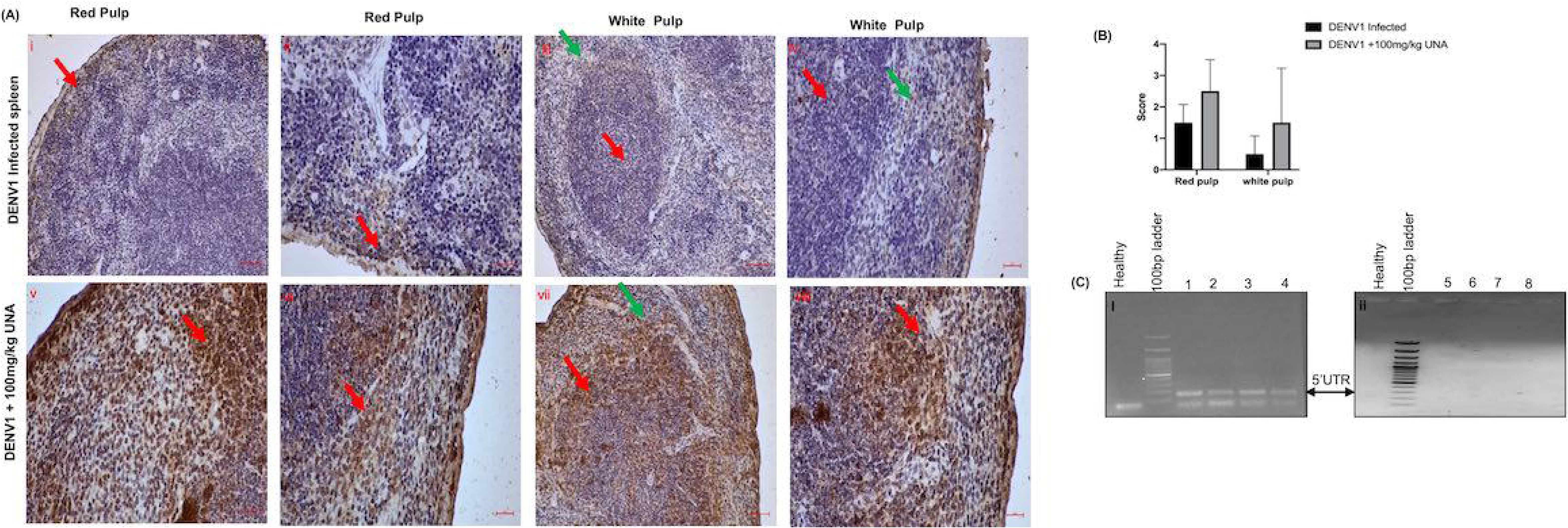
Immunohistochemical analysis of spleen tissues showing IFNGR1 expression in DENV1 infected and UNA-treated mice. (A) (i-ii) Mild cytoplasmic/membranous and nuclear expression of IFNGR1 protein in monocytes in red pulp region of spleen-(Red arrow); (iii-iv) Mild expression of cytoplasmic /membranous and nuclear expression of IFNGR1 protein in T lymphocytes in lymphoid follicles of white pulp-(Red arrow), mild expression in B lymphocytes in the marginal zone of white pulp bordering red pulp in the spleen-(green arrow); (v-vi) pronounced cytoplasmic /membranous and nuclear expression of IFNGR1 protein in monocytes in red pulp region of spleen-(Red arrow); (vii-viii) Pronounced expression of cytoplasmic/membranous and nuclear expression of IFNGR1 protein in T lymphocytes in lymphoid follicles of white pulp-(Red arrow), B lymphocytes in marginal zone of white pulp bordering red pulp in spleen-(green arrow). (B) Quantification of IFNGR1 expression. (C) RT-PCR analysis of viral RNA levels in blood samples collected on day 7 post-infection. Samples 1 to 4 represent infected mice while 5 to 8 represent UNA-treated infected mice(i-ii).

### SNTA1 binds to flavi and non-flavivirus capsid proteins

We performed a molecular docking study between SNTA1 and capsid proteins from both flaviviral (ZIKV, YFV,HCV and JEV) and non-flaviviral (CHIKV, ASFV and SARS-CoV-2) viruses using the ClusPro server. SNTA1 exhibited the strong binding with several flaviviridae members including ZIKV with a docking score of -1257.1, JEV (-1213.9), YFV (-1134.5) and moderate binding observed for HCV (-866.1) (Table S4). These interactions were supported by the presence of multiple hydrogen bonds and salt bridges, indicating stable complex formation. In contrast, non-flaviviral capsid proteins such as CHIKV showed weaker interactions (-794.3) while ASFV and SARS-CoV-2 also showed a significant score - 1171.3 and -1040.7 respectively (Table S6).

### Ursonic acid binds to the capsid proteins of flavi and non-flaviviridae members

Three-dimensional structures of capsid proteins from flaviviruses (ZIKV, YFV and JEV) and selected non-flaviviruses (CHIKV, ASFV, HCV and SARS-CoV-2) were modelled using the SWISS-MODEL server. The capsid structures were subsequently employed in molecular docking studies with UNA using AutoDock Vina. UNA exhibited the strong binding score with capsid/Nucleocapsid of several *flaviviridae* members including ZIKV showed a docking score of -12.0 kcal/mol, JEV (-10.3), YFV (-9.5) and moderate binding observed for HCV (-7.7) (Table S5). UNA showed high binding affinities towards non-flaviviral capsid proteins of SARS-CoV-2, which exhibited a strong interaction with binding energy: -11.2 kcal/mol, ASFV (-8.3) and CHIKV (-9.1) (Table S7). As SARS-CoV-2 N has shown strong binding with UNA so molecular dynamics simulations were performed for the SARS-CoV-2 nucleocapsid protein. Arg233, Lys217 formed hydrogen bonds with UNA. These interactions contributed to high conformational stability of the complex (Figures S6A-D). Further purified SARS-CoV2 nucleocapsid protein was used for fluorescence spectroscopy. The intrinsic fluorescence of the Nucleocapsid protein showed a decrease in fluorescence intensity upon interaction with increasing concentrations of UNA supporting the *in silico* analysis (Figures S6 A-G)

## Discussion

Mitochondria are proven to take part in resisting infections, especially in the form of innate immunity[51]. On the other side, viruses appear to have evolved to overcome the mitochondrial mediated resistance by applying possible mechanisms[8], [9], [52]. In this direction, dengue virus coded protease (non-structural protein-3; NS3) was shown to target the mitochondria and trigger them to the dysfunction[8], [9]. This protein exists in different forms (NS3 with and without NS2B) during infections and localizes to mitochondria and/or nucleus based on its association with NS2B[7], [8], [53]. Similarly, NS5 alone localizes to the nucleus but when associated with NS4B, it may be retained in the cytoplasm [12,53,54]. Recent studies have shown that HCV has three processed forms of capsid that localize to mitochondria and evade immune host response[12], [54], [55]. It was reported that the capsid protein of dengue virus localizes to the nucleus but the detailed mechanism is unclear. In light of the above literature, we carried out the time dependent DENV1 infection in the K562 cells and found four different capsid forms (c, c-anc and c-anc-pr, c-anc-prM) (Figure 1A). It appears that the polyprotein undergoes a systematic time dependent processing to yield the only capsid during the initial stages of the infection, that required for the formation of new viral particles at that stage. Then the polyprotein may switch its processing to yield the capsid along with its accessory proteins (c-anc, c-anc-pr and c-anc-prM) that are required to counter the host’s resistance during the later stages of the infection. We found the mature capsid in the nucleus and c-anc in mitochondria but the mitochondrial localization was especially strong for the c-anc and c-anc-pr form (Figure S1B and Figure 2A-E). This could be due to c-anc and c-anc-pr retain hydrophobic sequences, helping them stick to mitochondrial membranes just like the precursor forms of HCV’s core protein. To gain further deeper insights, we checked whether the c-anc localizes to inner or outer mitochondrial membranes. Results showed that it localizes to the outer membrane of mitochondria (Figure 2D and E). It appears that the capsid protein alone localizes mainly to the nucleus, while in association with anchor localizes to mitochondria. Given that many viruses are known to target mitochondria to modulate apoptosis, suppress immune responses and alter cellular metabolism, hence we asked, does this mitochondrial localization actually matter? We found that in the cells expressing the c-anc form, mitochondria became fragmented and their fusion proteins (MFN2 and OPA1) were downregulated suggesting that this protein alone can impair mitochondrial integrity (Figure 3A-F)[55]. Mitochondrial damage was shown earlier to be mediated by NS3 [8], [9], [56]. But, it appears that this virus may use multiple mechanisms to maximise mitochondrial damage. However, whether it is c-anc mediated or NS3 mediated in the initial order, or whether both occur together, is the subject of interest. Consistent with this, our data showed that key mitochondrial genes that involved in mitochondria function (ATP6, CO3, ND2 and CYB) were significantly downregulated in c-anc transfected cells (Figure S3A-C). Another interesting finding of the present study was the interaction between the c-anc and a host protein called alpha-1 syntrophin (SNTA1) (Figure S4A-C,4A). SNTA1 was consistently reduced in c-anc transfected, infected cells and patient serum samples and the more severe the infection, the lower the levels (Figure 4B-D). Previous studies have highlighted the role of SNTA1 in regulating oxidative stress and maintaining redox homeostasis. Overexpression of SNTA1 in H9C2 cells was shown to attenuate ROS accumulation, whereas SNTA1 knockdown led to elevated ROS levels, indicating its protective role against oxidative stress[57], [58], [59]. Furthermore, protein–protein interaction (PPI) network analysis further showed that SNTA1 interacts with key redox and cytoskeletal regulators, including MAPK12, a kinase known to modulate ROS responses, along with SNTG2, DMD, CAV3, and SGCA[58], [59]. These findings support the idea that SNTA1 contributes to cellular defence mechanisms against oxidative damage; hence, dengue virus may have targeted this protein to reduce its availability by c-anc mediated sequestering. Further analysis showed that mitochondria in c-anc expressing cells were malfunctioning: they had decreased mitochondrial potential, reduced respiration and less ATP (Figure 4E-G). In support of the above observations, dengue virus infected Vero cells have also shown decreased mitochondrial potential, reduced basal, maximal and non -mitochondrial respirations as well as ATP production and increased ROS (Figure 6E-G and S5E).

We made an attempt to identify a compound that inhibits c-anc, to counteract the above effects and ursonic acid (UNA) was identified in this direction (Figure 5A-D and Table 1). Computational modelling and fluorescence spectroscopy showed that UNA binds strongly to the dengue virus c-anc protein (Figure 5D-F). UNA inhibited virus multiplication in Vero cells and UNA helped to restore virus infected cells morphology and mitochondrial membrane potential as well as it decreases the ROS levels which are crucial for cell survival (Figures 6A-D; S5A-B; 6E ; S5E). Further, mitochondrial functions in infected cells were found to be normalized in the presence of this drug (Figure 6G [i-vi]). In addition, DENV1 infected C57BL/6 mice treated with UNA showed healthier spleen and better immune responses with the higher levels of IFNGR1 and reduced viral multiplication, possibly an immunomodulation effect of UNA (Figure 7 A-C). Through *in silico* analysis, we also found that capsid proteins from other flaviviruses like ZIKV, JEV and YFV interact with SNTA1, suggesting that this host interaction may be a conserved mechanism adopted by certain group of viruses (Table S4). Since mitochondrial damage is a common feature in infections caused by viruses such as HCV and SARS-CoV-2 and SNTA1, UNA or similar compounds could be used as potential anti-capsid molecules for the above virus infections.

### Quantification and stastical analysis

All quantitative data were expressed as mean ± standard deviation (SD) from at least three independent experiments. Statistical analyses were performed using GraphPad Prism 8. Student’s t-test, one-way ANOVA or two-way ANOVA were applied as appropriate based on the experimental design. Statistical significance was denoted as follows: p < 0.05 (*), p < 0.005 (**) and p < 0.00001(***) while "ns" indicates no significant difference. Densitometry of immunoblots was conducted using ImageJ (NIH). Fluorescence intensity quantification and mitochondrial morphology analyses were also performed using ImageJ with morphological parameters extracted using the Mitochondrial Analyzer plugin.

## Supporting information

all supplementary figures are referenced in manuscript

all supplementary Tables are referenced in manuscript

## Online Supplemental Material

Figure S1 shows the nuclear localization of dengue virus capsid and its variants in transfected cells. Figure S2 shows the localization of the capsid protein in infected cells. Figure S3 contains the transcriptomic profile of c-anc transfected cells. Figure S4 details the interaction between c-anc and SNTA1. Figure S5 demonstrates Ursonic acid decrease viral load, and improves infected cells morphology, mitochondrial potential, reduce ROS and improve hematology parameter in UNA-treated C57BL/6 mice. Figure S6 shows the interaction between Ursonic acid and the SARS-CoV-2 nucleocapsid. Table S1 lists the top 10 natural compounds with their ADMET properties. Table S2 and Table S6 provide protein-protein docking results for various viral capsids with SNTA1. Table S3, Table S5 and Table S7 provide molecular docking results for various viral capsids with ursonic acid.

## Data Availability

All data supporting the findings of this study are available within the article and its supplementary materials.

## IAEC and IEC approvals

Regulatory approvals for using the animals and human samples were obtained from the respective regulatory bodies of University of Hyderabad.

## Acknowledgements

The authors acknowledge the funding support from SERB, India (EMR/2016/007845), DBT, India (BT/PR14240/MED/29/985/2015) and DST-FIST level I, DOBB (SR/FST/LSI-530/2012). The authors also acknowledge the DBT-BUILDER program of School of Life Science (No BT/INF/22/SP41176/2020), UOH, India, for the Agilent Seahorse XFP mini analyzer and University Grants Commission (UGC) for providing fellowship to P.C.

