## Supplementary material for "Time-dependent processing of dengue virus polyprotein yields multiple capsid forms that disrupt cellular homeostasis": all supplementary figures are referenced in manuscript

**Supplemental Figures (S1 to S6)**

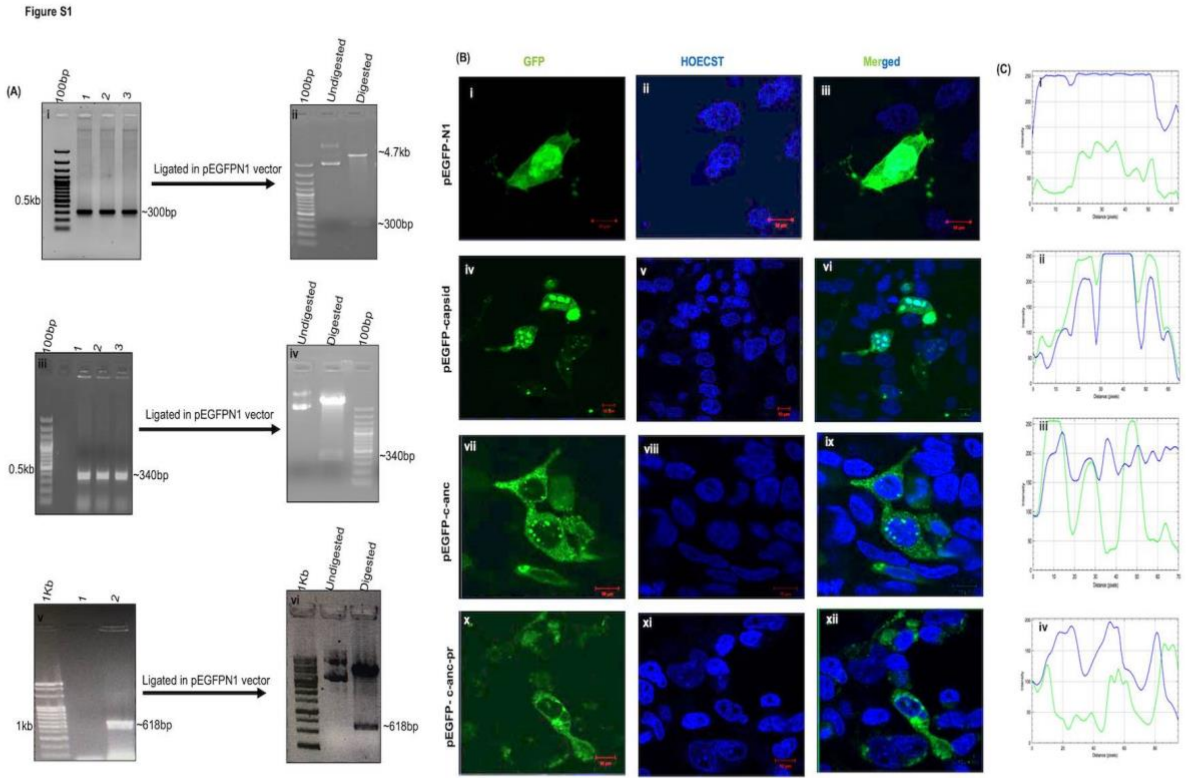


**Figure S1. Nuclear localization of dengue virus capsid and its variants in transfected HEK293 cells :(A)** Cloning of capsid variants in pEGFPN1 vector. Agarose gel electrophoresis showing PCR amplification of capsid variants and confirmation of clones by restriction digestion using *Xho* I and *Bam* HI: (i) lane1 showing 100bp ladder and 1,2,3 showing amplified capsid; (ii) restriction digestion of pEGFP-capsid ; (iii) lane 1 showing 100bp ladder and 1,2,3 showing amplified pEGFP-c-anc; (iv) restriction digestion of pEGFP-c-anc; (v) lane 1 showing 1kb ladder and 1,2 showing negative control and amplified pEGFP-c-anc-pr respectively; (vi) restriction digestion of pEGFP-c-anc-pr.(B) Confocal images acquired at 63× magnification show the localization of pEGFP-N1 vector (i–iii), pEGFP-c (iv-vi), pEGFP-c-anc (vii-ix), and pEGFP-c-anc-pr (x–xii), along with the nuclear localization marker, i.e., Hoechst 33342.(C) Fluorescence intensity profiles (i–iv) depict the distribution of GFP (green) and Hoechst (blue).

Figure S2

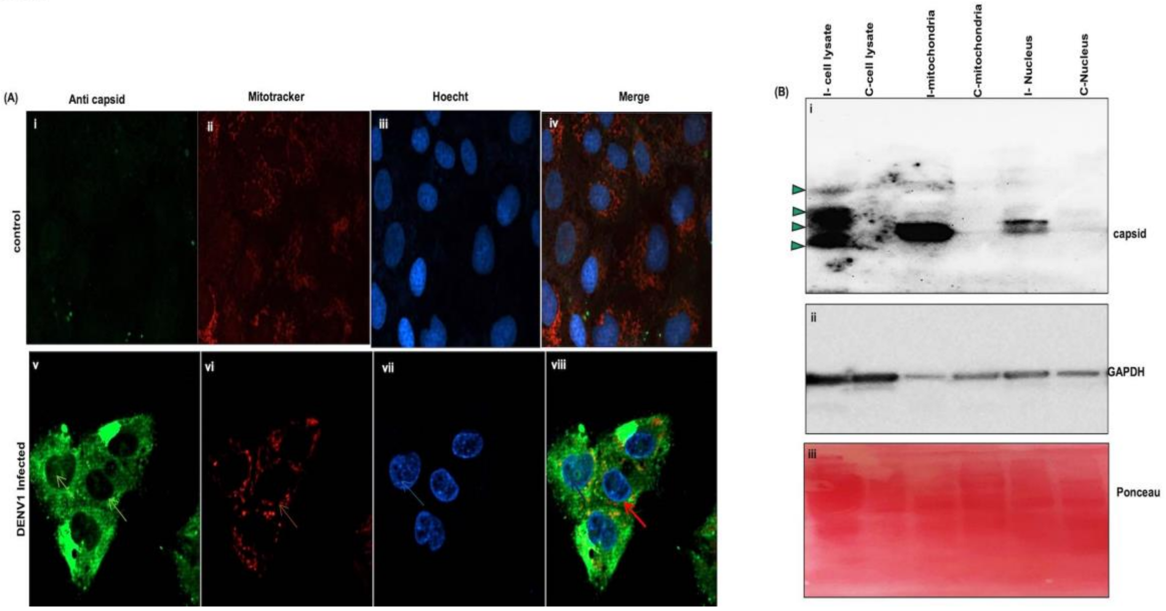


**Figure S2. Localization of dengue virus capsid protein in infected Vero cells**(A) Immunofluorescence images showing dengue virus capsid protein expression in Vero cells. Panels (i–iv): mock-infected cells; panels (v–viii): infected cells stained for nuclei (Hoechst), mitochondria (MitoTracker), and capsid protein (green). Blue arrows indicate nuclear and red arrow indicates mitochondrial localization of the capsid protein.(B) (i)Subcellular fraction of uninfected and dengue infected Vero cells; (ii) GAPDH (iii) Ponceau S stained membrane, probed and reprobed with the above antibodies. Arrow indicates the four forms of the capsid protein observed in infected cell lysate.

**Figure S3**

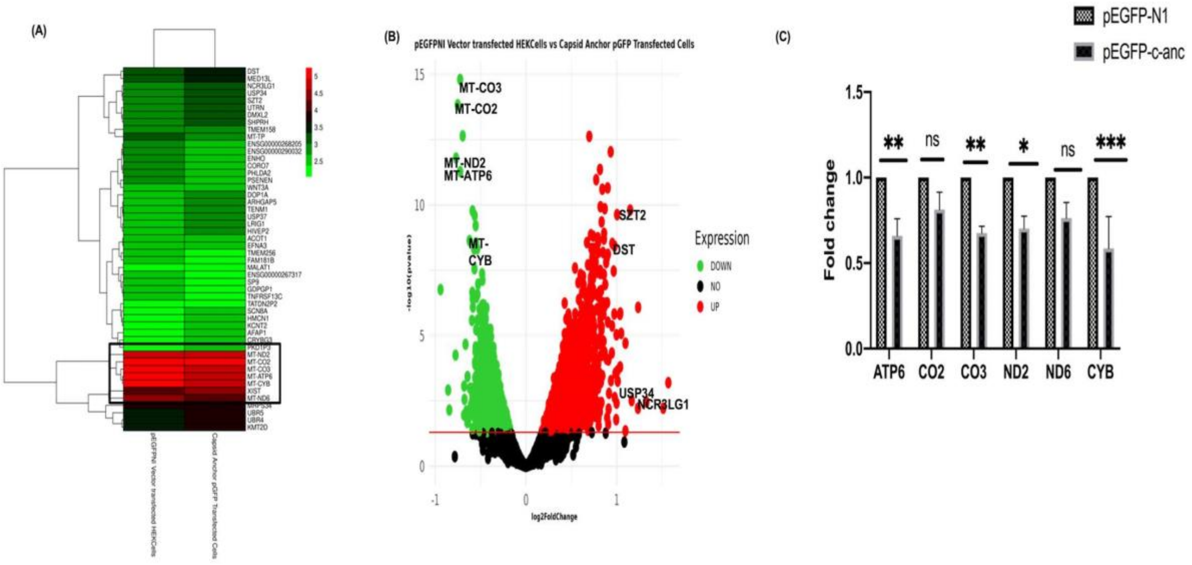


**Figure S3: Transcriptomic profiling in c-anc transfected HEK293 cells.**(A) Heatmap displaying the differentially expressed genes (DEGs) in pEGFP-c-anc transfected HEK293 cells compared to pEGFP-N1 transfected cells. (B) Volcano plot illustrating the distribution of DEGs. Red dots represent significantly upregulated genes and green dots represent significantly downregulated genes.(C) Quantitative real-time PCR (qRT-PCR) validation of selected mitochondrial genes in c-anc and pEGFP-N1 transfected cells. Bar graphs show relative mRNA expression normalized to GAPDH.

Figure S4

(A)

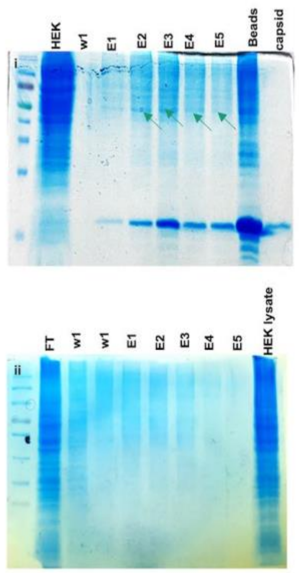


(B)

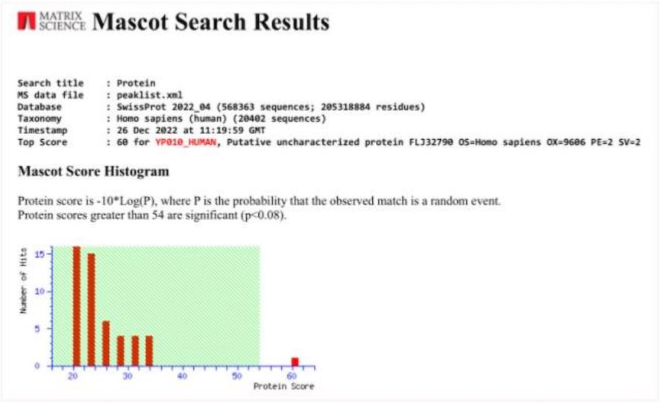


(C)

| | Accession | Mass | Score | Description |
|---|---|---|---|---|
| 1. | YP010_HUMAN | 21022 | 60 | Putative uncharacterized protein FLJ32790 OS=Homo sapiens OX=9606 PE=2 SV=2 |
| 2. | SNTA1_HUMAN | 54204 | 34 | Alpha-1-syntrophin OS=Homo sapiens OX=9606 GN=SNTA1 PE=1 SV=1 |
| 3. | KCD18_HUMAN | 47223 | 34 | BTB/POZ domain-containing protein KCTD18 OS=Homo sapiens OX=9606 GN=KCTD18 PE=2 SV=2 |
| 4. | DYN2_HUMAN | 98345 | 33 | Dynamin-2 OS=Homo sapiens OX=9606 GN=DNM2 PE=1 SV=2 |
| 5. | LACB2_HUMAN | 33070 | 33 | Endoribonuclease LACTB2 OS=Homo sapiens OX=9606 GN=LACTB2 PE=1 SV=2 |
| 6. | CCNB2_HUMAN | 45538 | 32 | G2/mitotic-specific cyclin-B2 OS=Homo sapiens OX=9606 GN=CCNB2 PE=1 SV=1 |
| 7. | SFSWA_HUMAN | 105157 | 31 | Splicing factor, suppressor of white-apricot homolog OS=Homo sapiens OX=9606 GN=SFSWAP PE=1 SV=3 |
| 8. | RBM45_HUMAN | 53868 | 31 | RNA-binding protein 45 OS=Homo sapiens OX=9606 GN=RBM45 PE=1 SV=1 |
| 9. | NDKM_HUMAN | 20703 | 31 | Nucleoside diphosphate kinase, mitochondrial OS=Homo sapiens OX=9606 GN=NME4 PE=1 SV=1 |
| 10. | SPC1L_HUMAN | 37932 | 29 | Speriolin-like protein OS=Homo sapiens OX=9606 GN=SPATC1L PE=1 SV=3 |
| 11. | YBEY_HUMAN | 19457 | 28 | Endoribonuclease YbeY OS=Homo sapiens OX=9606 GN=YBEY PE=1 SV=2 |
| 12. | PKHH2_HUMAN | 169720 | 28 | Pleckstrin homology domain-containing family H member 2 OS=Homo sapiens OX=9606 GN=PLEKHH2 PE=1 SV=2 |
| 13. | PTPRB_HUMAN | 225587 | 27 | Receptor-type tyrosine-protein phosphatase beta OS=Homo sapiens OX=9606 GN=PTPRB PE=1 SV=3 |
| 14. | TRAF7_HUMAN | 76386 | 27 | E3 ubiquitin-protein ligase TRAF7 OS=Homo sapiens OX=9606 GN=TRAF7 PE=1 SV=1 |
| 15. | ALG1L_HUMAN | 21575 | 27 | Putative glycosyltransferase ALG1-like OS=Homo sapiens OX=9606 GN=ALG1L PE=2 SV=2 |
| 16. | LRC20_HUMAN | 20553 | 26 | Leucine-rich repeat-containing protein 20 OS=Homo sapiens OX=9606 GN=LRRC20 PE=1 SV=1 |
| 17. | ZN577_HUMAN | 56101 | 26 | Zinc finger protein 577 OS=Homo sapiens OX=9606 GN=ZNF577 PE=2 SV=3 |
| 18. | TP4A1_HUMAN | 20144 | 26 | Protein tyrosine phosphatase type IVA 1 OS=Homo sapiens OX=9606 GN=PTP4A1 PE=1 SV=2 |
| 19. | PIRT_HUMAN | 15495 | 25 | Phosphoinositide-interacting protein OS=Homo sapiens OX=9606 GN=PIRT PE=2 SV=2 |
| 20. | ENPP3_HUMAN | 101998 | 24 | Ectonucleotide pyrophosphatase/phosphodiesterase family member 3 OS=Homo sapiens OX=9606 GN=ENPP3 PE=1 SV=2 |

**Figure S4: Interaction of c-anc with SNTA1.**(A) (i) In vitro pull-down assay using HEK293 cell lysate with c-anc purified protein; (ii) Negative control.(B) The graph represents the Mascot Search Result.(C) Proteins identified using MS/TOF Mascot search along with their accession numbers and Mascot scores.

**Figure S5**

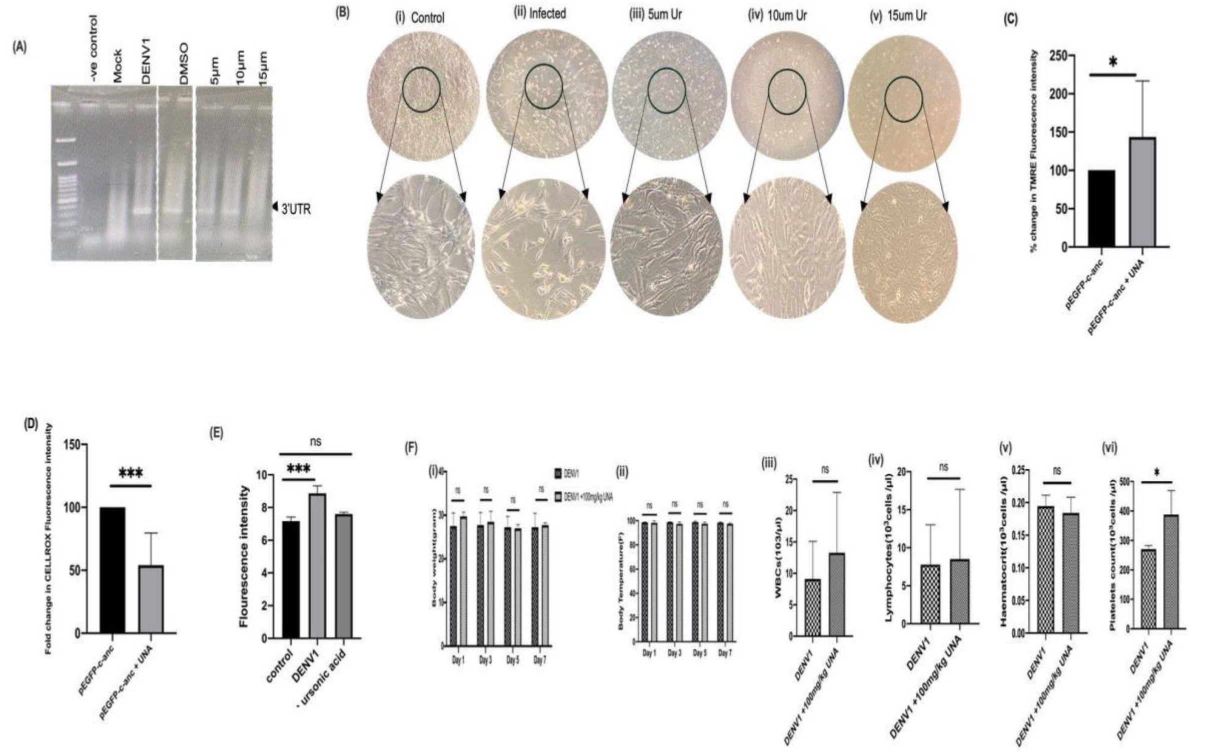


**Figure S5: Ursonic acid decrease viral load, and improves infected cells morphology, mitochondrial potential, reduce ROS and hematology analysis in UNA-treated C57BL/6 mice.**(A) Agarose gel electrophoresis image showing PCR analysis of supernatants from (i) DENV1 infected cells treated with UNA at 5, 10 and 15 µM concentrations. (B) Cell morphology analysis of Vero cells under DENV1-infected conditions following UNA treatment; (i–v) show uninfected, infected and UNA-treated cells at 5, 10, and 15 µM concentrations along with enlarged views of the corresponding cells.**(C**) Quantification of Mitochondrial membrane potential of c-anc transfected cells in presence and absence of UNA.(D) Quantification of ROS levels in pEGFP-c-anc transfected cells with and without UNA-treatment. (E) Measurement of cell ROS fluorescence intensity in uninfected, DENV1-infected and UNA-treated infected Vero cells using a CellROX™ Deep Red Reagent. (F) Hematology of infected and UNA-treated C57BL/6 mice; (i) Body weight; (ii) Temperature; (iii) WBCs; (iv) Lymphocyte; (v) Hematocrit; (vi) Platelet Count,

**Figure S6**

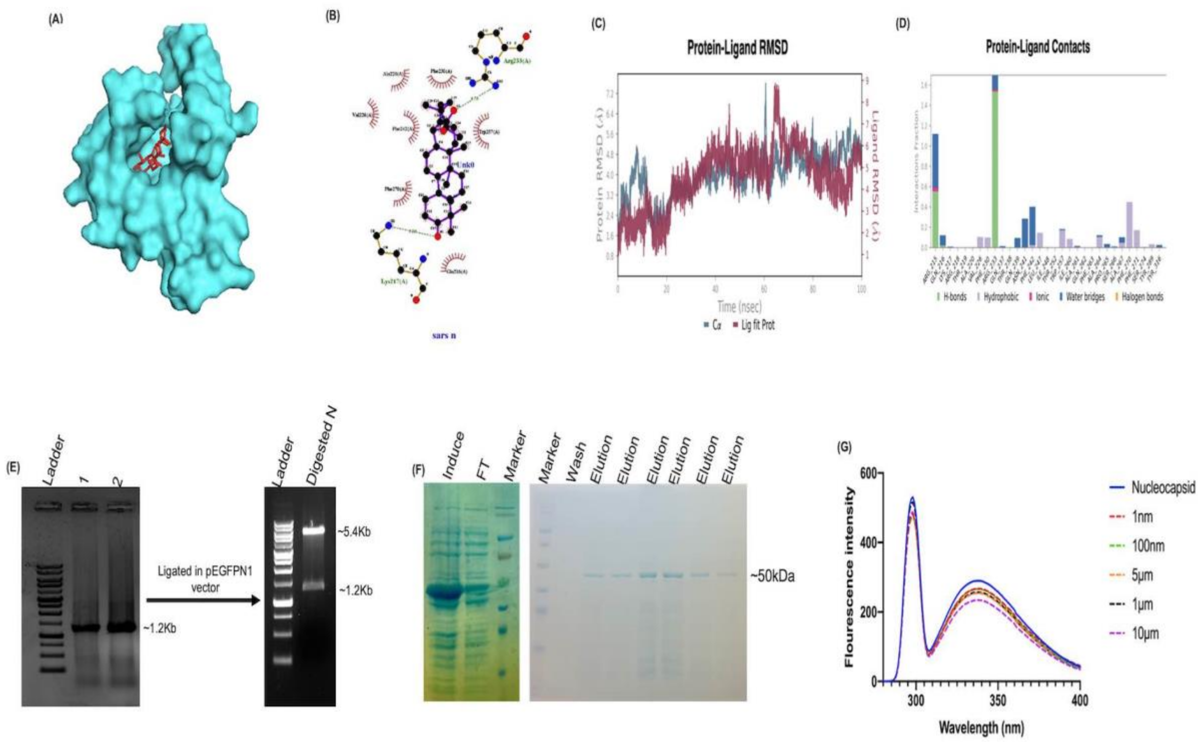


**Figure S6. Ursonic acid interact with SARS-CoV-2 nucleocapsid**(A) Molecular docking showing ligand UNA (stick model) binding to the SARS nucleocapsid protein.(B) Protein-ligand contacts showing the interacting residues, depicted by LigPlot. (C) RMSD analysis of the UNA bound nucleocapsid shows the changes in protein and protein-ligand complex stability over time during the dynamic simulation of the nucleocapsid.(D) Protein-ligand contacts interaction fraction calculated from the molecular dynamics simulation. (E) Cloning of SARS-CoV2 nucleocapsid in pET28a vector (i) lane1 showing 1kb ladder and 1,2 showing amplified nucleocapsid; (ii) confirmation of pET 28a-nucleocapsid by restriction digestion using *Eco* RI and *Bam* HI.(F) Expression and purification of the SARS-CoV-2 nucleocapsid protein; (i) The protein expression is shown in Lane 1, (ii)The protein ladder followed by washes and elution fractions.(F) Effect of ursonic acid on the intrinsic fluorescence of the nucleocapsid protein in the presence and absence of ursonic acid
