## Supplementary material for "Time-dependent processing of dengue virus polyprotein yields multiple capsid forms that disrupt cellular homeostasis": all supplementary Tables are referenced in manuscript

**Supplemental Tables (S1 to S7)**

| **Zinc ID** | **Molecular weight (MW)** | **H2 Bond Donors** | **H2 Bond Acceptors** | **Lipophilicity (clogp)** | **Giaa** | **Bbbpb** | **CYP1A2 inhibitor** | **CYP2C19 inhibitor** | **CYP2C9 inhibitor** | **CYP2D6 inhibitor** | **Molar Refractivity** |
|---|---|---|---|---|---|---|---|---|---|---|---|
| **ZINC13558221** | **454.68 g/mol** | **1** | **3** | **5.95** | **Low** | **No** | **No** | **No** | **No** | **No** | **135.95** |
| ZINC95914024 | 428.73 g/mol | 0 | 1 | 7.56 | Low | No | No | No | No | No | 135.88 |
| ZINC95486126 | 426.72 g/mol | 1 | 1 | 6.95 | Low | No | No | No | No | No | 135.40 |
| ZINC04721047 | 484.75 g/mol | 0 | 3 | 6.85 | Low | No | No | No | No | No | 144.06 |
| ZINC08551422 | 592.68 g/mol | 3 | 7 | 4.52 | Low | No | No | No | Yes | No | 180.42 |
| ZINC08951841 | 454.68 g/mol | 0 | 3 | 6.19 | Low | No | No | No | No | No | 133.56 |
| ZINC04721049 | 442.72 g/mol | 1 | 2 | 6.46 | High | No | Yes | No | No | No | 134.33 |
| ZINC95913853 | 440.66 g/mol | 1 | 3 | 6.05 | Low | No | No | No | Yes | No | 133.30 |
| ZINC02120777 | 501.57 g/mol | 2 | 6 | 4.82 | Low | No | No | Yes | Yes | No | 143.32 |
| ZINC08952018 | 472.70 g/mol | 3 | 4 | 5.23 | High | No | No | No | No | No | 137.82 |

**Table S1: Top 10 Natural Compounds with ADMET Properties and Lipinski's Rule of Five:** This table summarizes the Lipinski's Rule of Five and ADMET (Absorption, Distribution, Metabolism, Excretion and Toxicity) properties of the top 10 compounds selected from the virtual screening.

| S.No. | Capsid protein of Virus | Cluster Member | SALT Bridge | H2 Bond | Non Bonded Contacts | Score Docking | Interacting Residue | Pymol Visualization |
|---|---|---|---|---|---|---|---|---|
| 1 | DENV1 | 74 | 1 | 12 | 126 | -1144.8 | ALA46,ARG52, TRP66,LYS71, ARG95 | 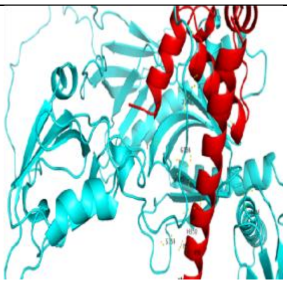 |
| 2 | DENV2 | 61 | 1 | 19 | 145 | -1086.3 | THR22,LEU41,LYS42,ALA49,LYS73,ASN76,LYS83, ARG87,ARG94,ARG96 | 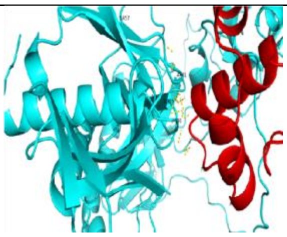 |
| 3 | DENV3 | 52 | 3 | 12 | 188 | -1077.9 | ASN21,ARG22,SER24,LYS45, ARG55,PHE56,LYS85,LYS86, ASN90,GLU87 | 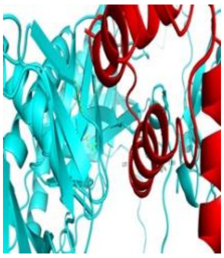 |
| 4 | DENV4 | 73 | 4 | 26 | 370 | -1086.3 | THR22,GLN24, LYS28,ARG29, THR32,ARG42, SER55,PRO57, PRO58,TRP66, LYS71,LEU92, ASN93,ARG95 | 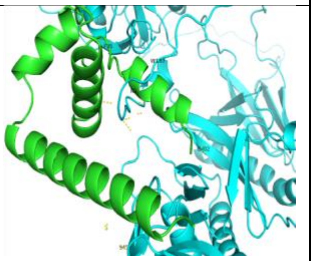 |

**Table S2: Protein–protein docking of capsid proteins of DENV1-4 with SNTA1 using ClusPro server**

| S.no. | Proteins | Interacting Residue (H2-bond) | Hydrophobic Interaction | Binding energy | Ligplot analysis |
|---|---|---|---|---|---|
| 1 | DENV2 | PHE53 | MET88,ILE85,PHE81,LEU78,ALA49,PHE50 | -11.6 | 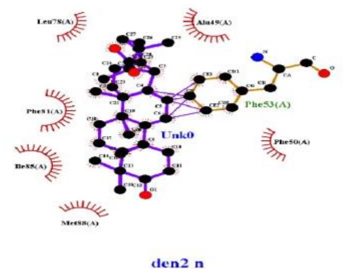  |
| 2 | DENV3 | ---- | ALA52,PHE53, MET91,ILE88,PHE84,ARG55,LEU81 | -12.1 | 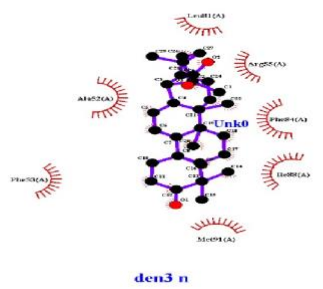  |
| 3 | DENV4 | ARG29 | PHE47,PHE50, PHE30,ALA46, LEU34,LEU26, MET43 | -11.5 | 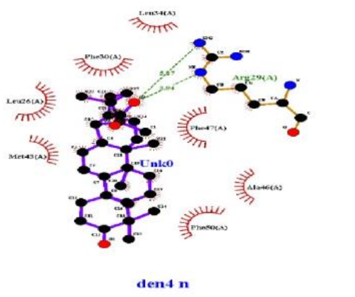  |

**Table S3. Molecular docking of dengue virus capsid proteins with ursonic acid using AutoDock Vina**

| S.No. | Capsid protein of Virus | Cluster Member | SALT BRIDGE | H2 BOND | NON BONDED CONTACTS | Score Docking | Interacting Residue | Pymol Visualization |
|---|---|---|---|---|---|---|---|---|
| **1** | ZIKV | 92 | 9 | 29 | 315 | -1257.1 | LYS18,ARG19, LEU27,LYS31, ARG32,LEU33, GLY36,HIS41, GLY42,ARG55, PRO61,LEU63, ARG68,SER71, LYS74,GLU76 ,LYS85 | 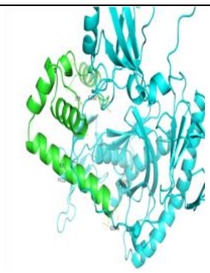 |
| **2** | YFV | 83 | 4 | 21 | 181 | -1134.5 | THR57,LYS60, ILE61,LYS66, ARG82,LYS85, ARG86,SER90, ARG93,ARG98 | 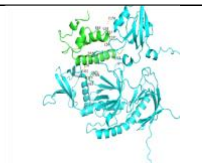 |
| **3** | JEV | 100 | 7 | 14 | 158 | -1213.9 | LEU27,GLY29, ARG32,ARG41, LYS55,LYS79, HIS80,LYS85, GLU87 | 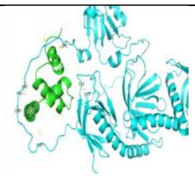 |
| 4 | HCV | 133 | 4 | 20 | 243 | -866.1 | ARG9,LYS12, ASN16,ARG18, ASP21,ARG39, PRO42,A RG43 | 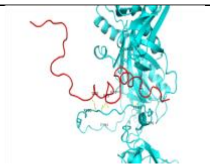 |

**Table S4: Protein–protein docking analysis of capsid proteins from other flaviviridae (ZIKV,YFV, JEV,HCV) with SNTA1**

| S.no. | Proteins | Interacting Residue (H2-bond) | Hydrophobic Interaction | Binding energy | Ligplot analysis |
|---|---|---|---|---|---|
| 1 | ZIKV | PHE56, | ALA49,ALA52, PHE53,PHE84,ASP 87,MET91,LEU 88 | -12 | 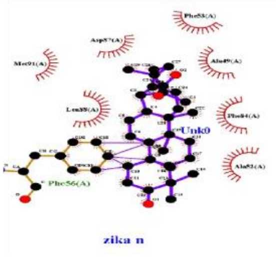 |
| 2 | YFV | ---- | PHE50,49,53,L EU56,52,LEU5 2,56,VAL84,GL Y46 | -9.5 | 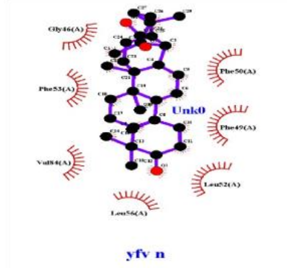 |
| 3 | JEV | PHE56, ARG45, | THR52,LEU88, GLU87,PHE84, ALA49 | -10.3 | 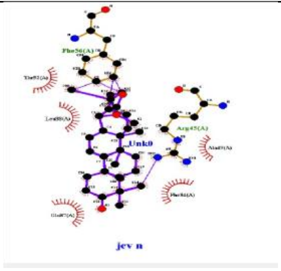 |
| 4 | HCV | ASN16,AS N14,THR15 ,GLN8 | PRO19,LYS12, ARG9,ARG13 | -7.7 | 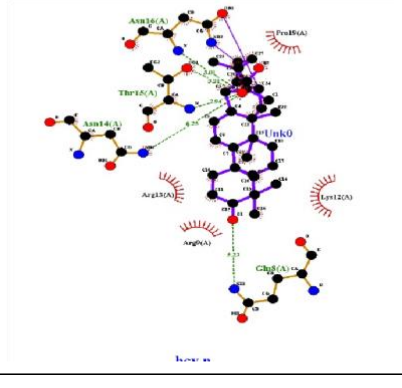 |

**Table S5. Molecular docking of capsid proteins from other flaviviridae (ZIKV,YFV, JEV,HCV) with ursonic acid using AutoDock Vina**

| S.No. | Capsid protein of Virus | Cluster Member | SALT BRIDGE | H2 BOND | NON BONDED CONTACTS | Score Docking | Interacting Residue | Pymol Visualization |
|---|---|---|---|---|---|---|---|---|
| **1** | CHIKV | 67 | 2 | 18 | 187 | -794.3 | LYS51,ARG52, SER53,LYS55, TYR56,LYS143, LYS148, GLY153, ALA154, GLU156 | 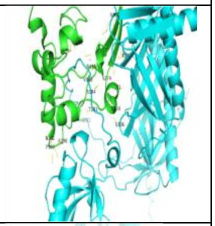 |
| 2 | ASFV | 71 | 1 | 19 | 201 | -1171.3 | ASN53,GLN57, HIS66,LYS67, VAL71,ASN73, SER101,SER104, PRO109,ASP191, ASP153,TYR192 | 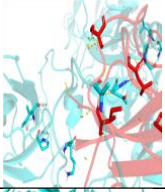 |
| 3 | SARS-CoV-2 | 91 | 4 | 23 | 283 | -1040.7 | LYS213,ARG215, LYS217,PRO235, GLU236,GLN237, THR238,PHE271, MET273,SER274, ARG275,GLU279 , THR285,LYS317 | 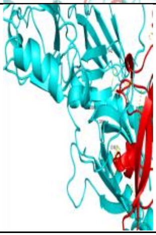 |

**Table S6: Protein–protein docking analysis of capsid/Nucleocapsid proteins from non-flaviviridae with SNTA1**

| S.no. | Proteins | Interacting Residue (h-bond) | Hydrophobic Interaction | Binding energy | Ligplot analysis |
|---|---|---|---|---|---|
| 1 | CHIKV | ASP144,HIS76,TRP141,SER53 | LYS29,LEU58, CYS16,TRP56, VAL46,PHE74, ASP28 | -9.1 | 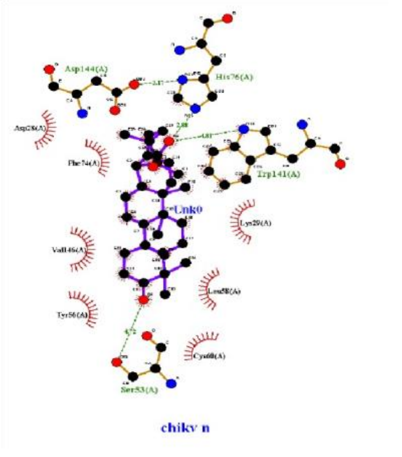 |
| 2 | ASFV | LYS150 | PRO152,ALA157,GLY47,ISO46,SER204,PHE68,PH45,VAL202 | -8.3 | 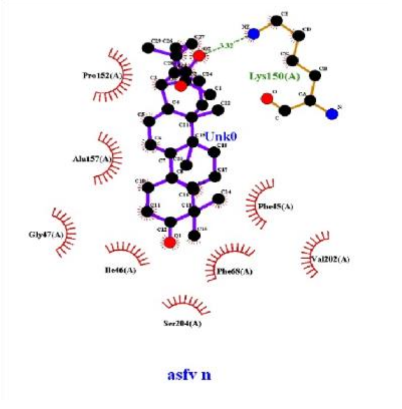 |
| 3 | SARS-CoV-2 | ARG233,LYS217 | TRP257,GLN216,PHE270,PHE242,PHE230, ALA220,VAL226, | -11.2 | 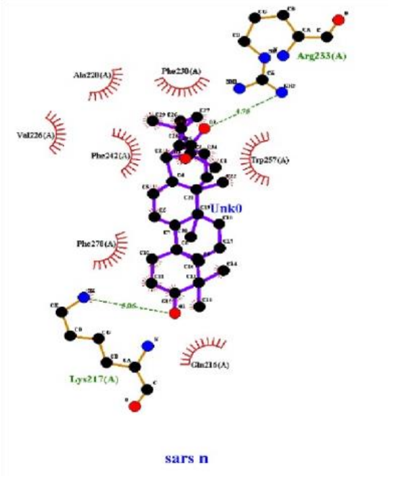 |

**Table S7. Molecular docking of capsid/Nucleocapsid proteins from non-flavivirus with ursonic acid using AutoDock Vina**
